# Amodal population clock in the primate medial premotor system for rhythmic tapping

**DOI:** 10.1101/2022.08.14.503904

**Authors:** Abraham Betancourt, Oswaldo Pérez, Jorge Gámez, Germán Mendoza, Hugo Merchant

## Abstract

The neural substrate for beat extraction and response entrainment to auditory and visual rhythms is still unknown. Here we analyzed the population activity of hundreds of medial premotor neurons of monkeys performing an isochronous tapping guided by brief flashing stimuli or auditory tones. The animals showed a strong bias towards visual than auditory metronomes, with rhythmic tapping that was more precise and accurate on the former. The population dynamics shared the following properties across modalities: the circular dynamics of the neural trajectories formed a regenerating loop for every produced interval; the trajectories converged in similar state space at tapping times resetting the clock; the tempo of the synchronized tapping was encoded in the trajectories by a combination of amplitude modulation and temporal scaling. In addition, the modality induced a displacement in the neural trajectories in auditory and visual subspaces without greatly altering time keeping mechanism. These results suggest that the interaction between the MPC amodal internal representation of pulse and a modality specific external input generates a neural rhythmic clock whose dynamics governs rhythmic tapping execution across senses.

## Introduction

Humans have the natural ability to extract salient periodic events from sound sequences, called the beat or pulse, and to predictively align their movements to this pulse in music and dance (Honing, 2012; Merchant et al., 2015b). Importantly, these cognitive abilities depend on an internal brain representation of pulse that involves the generation of regular temporal expectations (Balasubramaniam et al., 2021). Many studies have shown that the internal pulse is directly mapped to the timing of the entrained movements, typically measured as tapping or head bopping movements that occur few milliseconds before the beat (Lenc et al., 2021; Nozaradan et al., 2016, 2017). In addition, functional imaging in humans have suggested that the neural substrate of the internally-represented pulse lies in the motor system, including the medial premotor areas (MPC: SMA and pre-SMA) and the basal ganglia (Grahn & Rowe, 2009; Kung et al., 2013; Sánchez-Moncada et al., 2020). Influential ideas suggest that beat perception and entrainment depend on the periodic activation of the motor control regions of the brain that simulate regular actions in a predictive fashion (Cannon & Patel, 2021; Merchant & Yarrow, 2016; Patel & Iversen, 2014). In addition, a crucial component of rhythmic entrainment is the error correction in the tapping sequence, which allows for the compensation of movement timing to do not lose the series of pulses. This compensation includes a conscious awareness and depends on an internal keeper mechanism that controls the period between taps, so that an interval that is short tends to be followed by a long interval and vice versa, avoiding large error accumulation (Jantzen et al., 2018; Repp & Keller, 2004).

The amazing human flexibility for perceiving and entraining to complex musical pieces with a sophisticated metrical structure seems to be species-specific (Fitch, 2013). Nevertheless, behavioral and EEG studies in macaques have strongly suggested that monkeys possess all the audiomotor machinery to perceive and entrain to isochronous metronomes (Honing, 2012; Honing et al., 2018; Zarco et al., 2009). Indeed, monkeys generate tapping movements in anticipation of the metronome and can flexibly change their movement tempo from trial to trial covering a range from 400 to 1000ms (Gámez et al., 2018; see also García-Garibay et al., 2016; Takeya et al., 2017). Furthermore, monkeys can superimpose accentuation patterns onto an isochronous auditory sequence, suggesting that they have the ability to generate a simple subjective rhythm on a regular auditory sequence (Ayala et al., 2017; Criscuolo et al., 2021). These findings support the gradual audiomotor hypothesis that suggests that beat-based timing emerged gradually in primates, peaking in humans due to a sophisticated audiomotor circuit, but present also for isochrony in macaques because of the close interaction between MPC, the basal ganglia, and the auditory cortex (Merchant & Honing, 2014). Indeed, recent neurophysiological studies in monkeys indicate that the internal pulse representation during rhythmic tapping to visual metronomes depends on the neural population dynamics in MPC. A key property of MPC neurons is the relative representation of beat timing. Cells that encode elapsed or remaining time for a tap show up-down ramping profiles that span the produced interval, scaling in speed as a function of beat (Merchant et al., 2011; Merchant & Averbeck, 2017). In addition, these cells are recruited in rapid succession producing a progressive neural pattern of activation (called neural sequences or moving bumps) that flexibly fills the beat duration depending on the tapping tempo, providing a relative representation of how far an interval has evolved (Crowe et al., 2014; Merchant et al., 2015a; Zhou et al., 2020). Another critical aspect of the MPC beat-based clock is that it resets on every interval providing an internal representation of pulse (Merchant et al., 2015b; Merchant & Bartolo, 2018). The neural cyclic evolution and resetting are more evident when the time-varying activity of MPC neurons is projected into a low-dimensional state space (Gámez et al., 2019). The population neural trajectories show the following properties. First, they have circular dynamics that form a regenerating loop for every produced interval. Second, they converge in similar state space at tapping times, resetting the beat-based clock at this point. Finally, the periodic trajectories increase in amplitude as a function of the length of the isochronous beat (Gámez et al., 2019; Lenc et al., 2021).

A fundamental unanswered question is whether the internal beat representation in MPC works as a general clock across metronome modalities or whether visual and auditory periodic stimuli engage MPC neural populations with temporal processing dynamics that are modality specific. The classical notion of a common clock across timing contexts (Buhusi & Meck, 2005; Gibbon et al., 1984; Treisman, 1963) has been replaced by the hypothesis of a neural timing mechanism that includes both a core timing network rooted on the motor system and a set of areas that are selectively engaged depending on the specific requirements of a task (Coull et al., 2011; Merchant et al., 2013a; Wiener et al., 2010). If the former notion is true, we should expect that the neural trajectories in MPC should have very similar properties when monkeys execute rhythmic tapping sequences in synchrony with auditory or visual metronomes. In contrast, if the later prevails we should observe neural population dynamics with some shared properties but also temporal processing that is modality specific.

To address these possibilities, we recorded the simultaneous activity of MPC cell populations while monkeys performed a synchronization tapping task guided by brief isochronous flashing stimuli or auditory tones. The animals showed a strong bias towards visual metronomes, with rhythmic tapping that was more precise and accurate than with auditory metronomes. A detailed analysis on the neural population trajectories and on the neural patterns of activation showed the following shared properties across modalities: (1) the circular dynamics of the neural trajectories and the neural sequences form a regenerating loop for every produced interval, producing a relative time representation; (2) the trajectories converge in similar state space at tapping times while the moving bumps restart at this point, resetting the beat-based clock; (3) the tempo of the synchronized tapping is encoded by a combination of amplitude modulation and temporal scaling in the neural trajectories, which at the level of neural sequences relates with a mixture of an increase in the number of engaged neurons, larger recruitment lapses between neurons, and a rise in the duration of their activation periods. On the other hand, the modality induced a large displacement of the neural trajectories without greatly altering their cyclical organization, duration dependent changes in amplitude and temporal scaling, nor the tap separatrix behavior. These later results are in line with the notion of a modality dependent tonic external input that produced a divergence in the cyclic neural trajectories to different subspaces. Therefore, these results suggest that the interaction between the amodal internal representation of pulse within MPC and a modality specific external input generates a neural rhythmic clock whose dynamics produce the differences in temporal execution between the auditory and visual metronomes.

## Results

Two monkeys were trained in a synchronization tapping task (ST). The animal started by placing his free hand on a key, kept it there for two stimuli (beat perception epoch), and then tapped a push button in response to the isochronous stimuli, producing five rhythmic intervals in a sequence (synchronization epoch; Figure 1A. See Methods). Brief auditory or visual stimuli were used as metronomes with an interonset interval of 450 or 850 ms in blocks of 25 trials. The order of the four interval/modality combinations was random across days. We are assuming that monkeys solved this task by generating an internal representation of the metronome’s pulse that is coupled one-to-one with the tapping times. This coupling has been extensively demonstrated in humans and seem to generalize in macaques. In addition, an error correction mechanism should be in place to maintain tap synchronization with the metronome, since in humans a longer produced interval tends to be followed by a shorter interval, while a shorter interval tends to be followed by a longer produced duration(Iversen et al., 2015).

**Figure 1.**
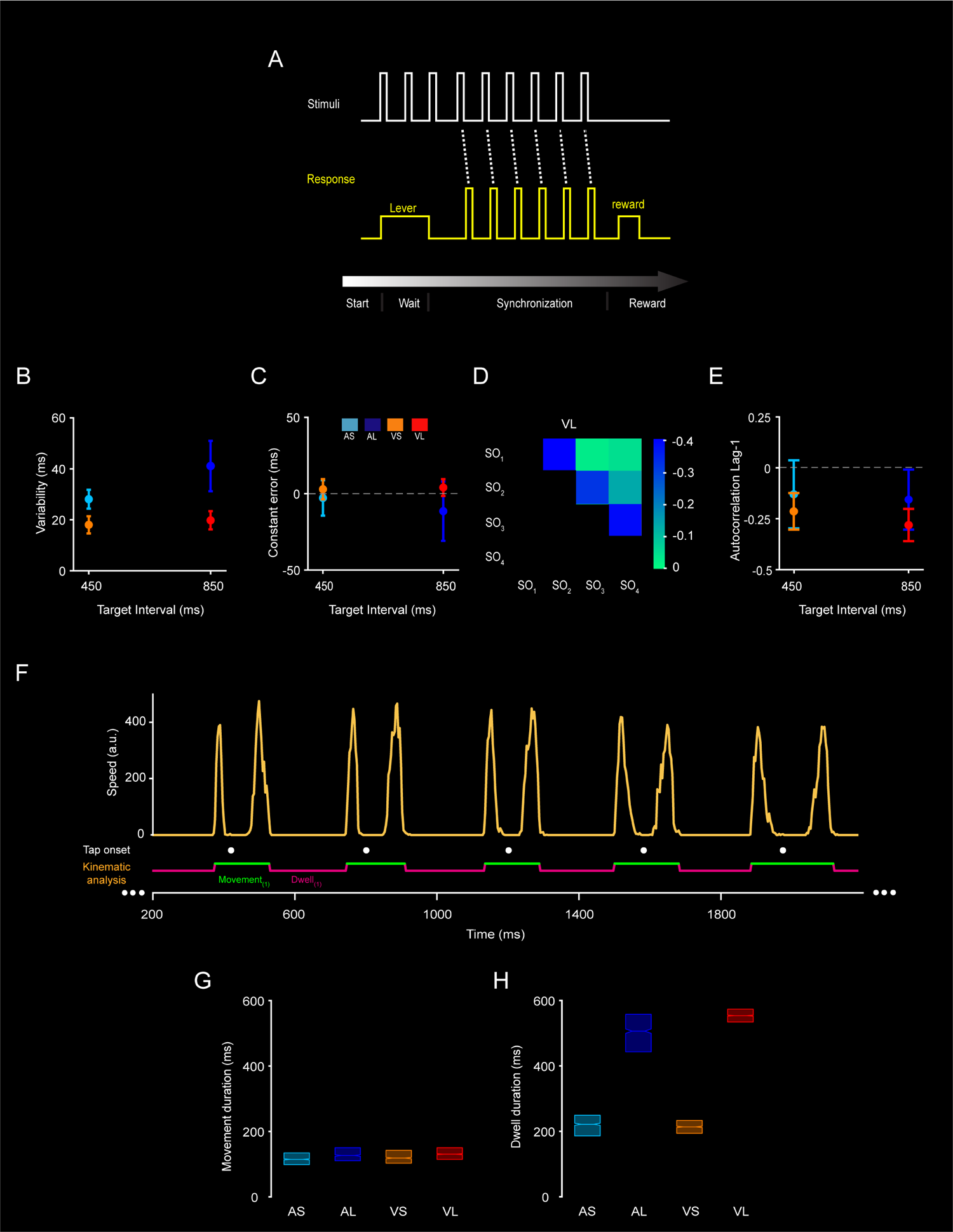
Rhythmic tapping behavior. (A) Synchronization task (ST). A trial started when the monkey placed his hand on a lever for a variable delay period. Then, a visual or auditory metronome was presented, and the monkey tapped on a button to produce five intervals following the isochronous stimuli of a particular duration. Correct trials were rewarded with an amount of juice that was proportional to the trial length. The instructed target durations were 450 and 850 ms. The first produced interval was not used for behavioral and neurophysiological analysis, since includes the movement from the key to the button. (B) Temporal variability (± 5xSEM) for each instructed interval and metronome modality, respectively (A: auditory, blue, V: visual, red). Shaded regions indicate ± ½ standard deviation from the mean. (C) Constant error (± 5xSEM) as a function of target interval during the auditory and visual condition of the ST. Inset: boxes indicate the ST condition, Cyan: 450 ms short auditory (AS); Blue: 850 ms large auditory (AL); Orange: 450 ms short visual (VS) and Red: 850 ms large visual (VL). (C) The lag 1 autocorrelation (± 5xSEM) for each instructed interval and metronome modality, respectively. (D) Correlation matrix of the produced intervals in the sequence (from serial order [SO] 1 to 4) within trials for the target duration of 850 ms in the visual condition. Note the large negative correlation for consecutive intervals. (F) Movement kinematics during ST. Speed profile of the hand movement (yellow trace) for the 450 ms interval of the auditory condition from the first to the sixth tap of a typical trail. Tap times are represented as white dots. The monkey produced highly stereotypical tapping movements with a constant duration flanked by dwell periods whose duration scaled with the metronome’s tempo. (G-H) Box plots (median and interquartile values) for the movement and dwell times, respectively, for each instructed interval and metronome modality.

We used the constant error (the difference between produced and instructed durations) and temporal variability (SD of produced intervals) to determine the accuracy and precision of the internal representation of the pulse based on the tapping times (Gámez et al., 2018; Zarco et al., 2009). A repeated-measures ANOVA on constant error revealed significant main effects for modality (F(1,87) = 15.57, P = 0.0002) and for the duration-modality interaction (F(1,87) = 4.1, p = 0.0457) (Figure 1C), but no duration (F(1,87) = 2.33, p = 0.13). This indicates that even when monkeys accurately produced intervals with errors close to zero, they slightly underestimated the intervals in the auditory condition, especially for the longer interval. The corresponding ANOVA on temporal variability showed significant main effects of target duration (F(1,87) = 39.53, p < 0.0001) and modality (F(1,87) = 146.49, p < 0.0001), as well as a significant effect for the duration x modality interaction (F(1,87) = 21.85, p < 0.0001) (Figure 1B). A Tukey honest significant difference (HSD) post hoc test showed a significantly larger temporal variability in the auditory than the visual condition, accompanied by an increase in slope in the temporal variability as a function of interval. These results indicate that timing precision in monkeys follows the scalar property of timing with a larger slope for auditory metronomes.

Figure 1D shows a large negative correlation for consecutively produce intervals (from serial order 1 to 4) for the target duration of 850 ms in the visual condition, indicating an error correction mechanism. Hence, we measured the autocorrelation of the inter-tap interval time series within trials and focused on its magnitude at lag 1. The lag 1 autocorrelation (Figure 1E) showed significant differences between modalities (F(1,87) = 15.08, p = 0.0009), and a marginal difference for target duration (F(1,87) = 3.4426, p = 0.078), but no statistical significant effect on their interaction (F(1,87) = 0.99, p = 0.33). Hence, these findings provide evidence of a stronger error correction mechanism for visual than auditory metronomes.

We further characterized the monkeys’ behavior during the ST by measuring the tapping movement kinematics. The speed profile of the hand movements captured by a high-speed camera was computed to determine the movement onset, the duration of each tapping, as well as the dwell time between tapping movements in monkey M2 (see Methods; Figure 1F). As previously observed in other animals during tapping tasks (Donnet et al., 2014; Gámez et al., 2018), the monkey showed a phasic stereotypic movement to push the button during ST (Figure 1F), with two bell-shaped speed kinematics (corresponding to the downward and upward movements, respectively) divided by a low-speed point associated with the button press. Thus, using a speed threshold we reliably identified four movement and four dwell times for each trial of our task (Figure 1F). Consistent with the notion of tapping stereotypy, the movement time was similar across the four conditions (Figure 1G). In contrast, the dwell time showed a large difference between shorter and longer target durations (Figure 1H). We carried out an ANOVA using the behavioral time dependent parameter and epoch (movement vs dwell time), duration and modality as factors, which showed significant main effects of epoch (F(1,13) = 502.51, p < 0.0001), duration (F(1,13) = 4560, p < 0.0001), and modality (F(1,13) = 20.12, p < 0.0001), and a large duration x epoch interaction (F(1,13) = 1910, p < 0.0001). Indeed, the post hoc Tukey HSD showed a large increase in dwell time between the target durations of 450 and 850 ms for both modalities (p < 0.0001).

Overall, these findings support the idea that during ST the monkeys used a complex rhythmic timing mechanism that includes at least four components. One for controlling the dwell to follow the metronome with different tempos (Figure 1I), another for triggering the internal beat signal that coincides with the tapping times, a component that generates a command that initiates the stereotypic 2-element tapping movement, and a mechanism for error correction within the rhythmic sequence. Furthermore, the modality of the metronome produced profound changes in rhythmic behavior, with timing that is more precise, more accurate, and with tap synchronization that showed larger error correction when using visual rather than auditory isochronous stimuli.

## Neural Trajectories

### General properties

The time-varying activity of a population of 1019 MPC neurons that fulfilled the criteria of number of trials and strength of responses during the execution of ST (see Methods) was projected into a low-dimensional space using principal component analysis (PCA) or Gaussian Process Factor Analysis (GPFA) (see Methods).

The resulting neural trajectories in state space showed periodic dynamics that formed an elliptical regenerating loop for every produced interval with properties that were dependent on time encoding, target duration, modality, and serial order of the produced interval within the sequence of ST (Figure 2A). We characterized these properties using geometric and kinematic approaches. Regarding the geometry, we first defined the subspaces for time encoding and each of remaining three task parameters by projecting the neural trajectories into second level PCs (PCs) over the 3D neural trajectories of the parameter of interest. For example, the plane for the auditory modality was defined by projecting the data into the first two second level PCs of the auditory neural trajectories in Figure 2A (Blue and Cyan trajectories).

**Figure 2.**
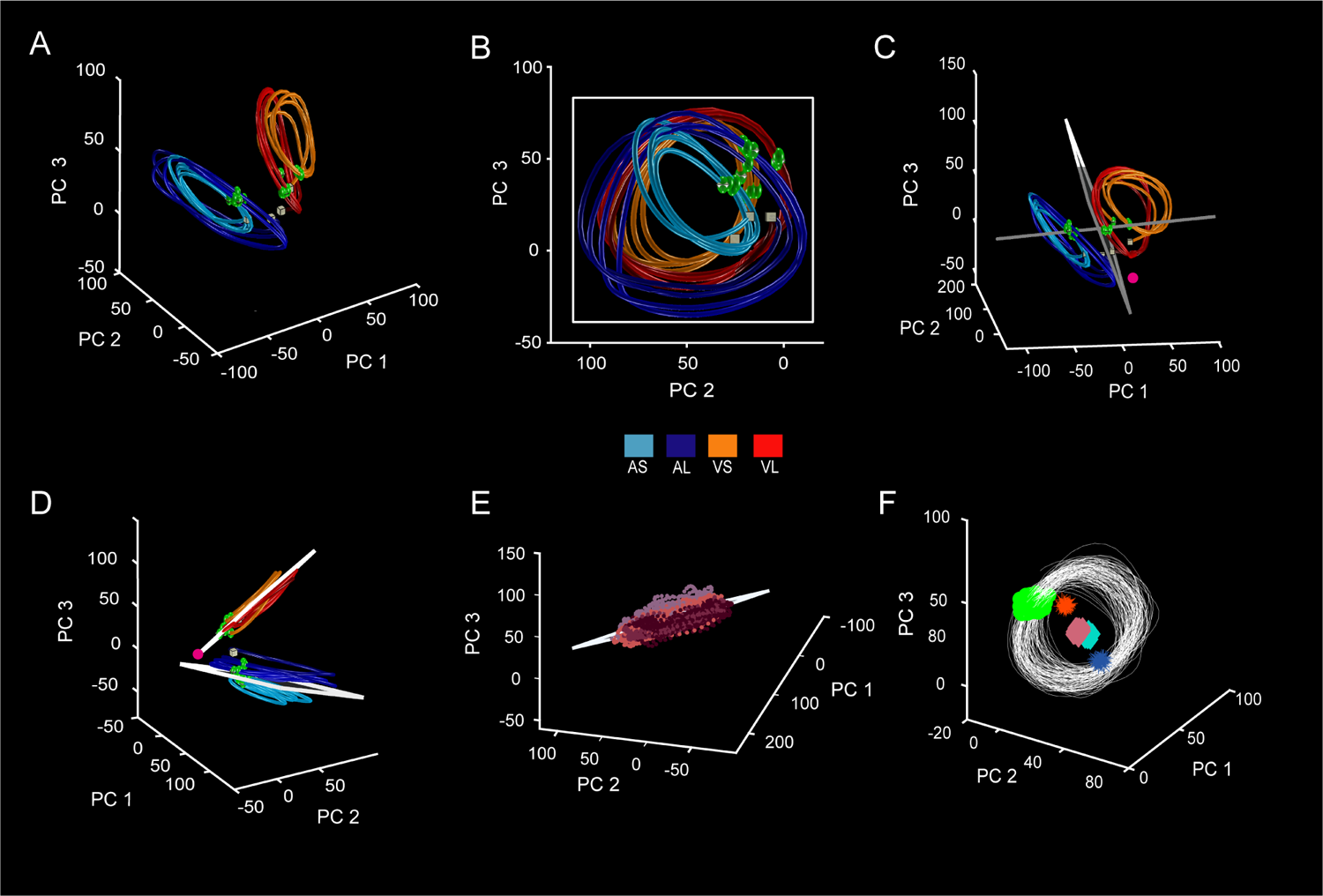
Neural population trajectories during ST and their oscillatory dynamic properties. (A) Projection of the neural activity in MPC (1,019 neurons) during ST onto the first three PCs. The first three PCs explained the 7.1, 4.1, and 4% of the total variance. Each point in the trajectory represents the neural network state at a particular moment, where the trajectory completes an oscillatory cycle on every produced interval across conditions. Cyan: 450 ms auditory; Blue: 850 ms auditory; Orange: 450 ms visual; Red: 850 ms visual. The green spheres indicate the tapping times across the trial sequence. (B) Neural population trajectories projected into duration plane, which explained the 45.8, 28.11, 25.9% of the variance for each PC. Target interval in milliseconds is color coded. A cube indicates the beginning of each trajectory, and an octahedron indicates the end. (C) Projection of the neural activity in the tapping time subspace, where the first three PCs explained 59.2,26.3, and 14.3% of the variance. (D) Projection of the neural activity in the modality subspaces. The first three PCs explained the 58.4, 38.9, and 2.5% of the variance for Auditory plane, while 58.3, 36.5, and 5% for visual plane. (E) Projection of the neural activity in the serial order subspace for VS condition explained 57.5%, 40.7% and 1.7% of the variance. Color code in B, C and D, same as in A. (F) General codification of the passage of time across the four conditions (white trajectories), as well as the overall state space values for target duration (cyan and cinnamon diamonds), modality (red/blue asterisks) and tapping times (green spheres).

Thus, the coding subspace for duration over time illustrated in Figure 2B shows periodic neural trajectories whose amplitude increase for longer target durations. This subspace (a plane defined by PCs 2 and 3) explained 96.7% of the variance of the trajectories and suggest the existence of a structured bimodal mechanism for tempo tracking where the amplitude of the rotatory neural dynamics defines the time between tapping movements (Figure 2B). In addition, the neural trajectories showed some degree of temporal scaling, stretching for short and compressing for long target intervals, with a scaling index of 0.88, 0.72 and 0.78 for the first three PCs of the auditory condition, and 0.86, 0.71 and 0.72 for the visual condition. Fully scaled trajectories should show a scaling index of one (Figure S3). Hence, time encoding depends on a mixed strategy that combines changes in the amplitude and speed of the neural trajectories. Notably, the population neural trajectories also converged in similar state space at tapping times, forming a line separatrix or boundary in state space was determined as the first PC of the tap locations in the trajectories across all serial order loops of the four conditions (Figure 2C). We hypothesize that a tick of the internal pulse representation was generated every time that the neural trajectories reached this separatrix. These findings imply that MPC is encoding three aspects of rhythmic timing: (1) the intertap interval is defined by each neural trajectory loop forming a limit cycle, where one periodic revolution is linked to a produced interval irrespective of its duration, encoding time in relative rather than absolute terms; (2) the target duration of the metronome using both a change in amplitude and temporal scaling for each cycle in state space; (3) the tapping time at the neural separatrix. Importantly, the angle between the duration subspace and the tap separatrix line was 88.3 degrees, suggesting an orthogonal arrangement where the internal pulse representation is independent of the time tracking neurons.

On the other hand, the trajectories for modality over time defined distinctive planes for the auditory and visual conditions explaining 97.5% and 94.9% the variance of the trajectories, respectively (first 3 PCs, Figure 2D). These subspaces had 54.6 degrees difference between modalities, backing the idea of a partially overlapping audiomotor processing in MPC (Figure 2D). We also found that the angle between duration and auditory subspaces was 24.3^*◦*^ and between duration and visual plane was 30.4°. These results indicate that the duration and modality subspaces are far from orthogonal and, hence, use similar but not identical neural resources. This is consistent with the mixed selectivity of single cells reported below. Finally, the serial order in the trial sequence has little influence on the neural trajectories, generating a tin subspace whose variability is below 2.3% (Figure 2E right), suggesting a small impact of serial order in rhythmic timing processing. We also used the Gaussian Process Factor Analysis (GPFA) as an alternative dimensional reduction technique that puts emphasis on the time series evolution rather than on the neural covariance of PCA. The neural trajectory properties were similar using PCA and GPFA (Figure S1), suggesting that they are not dependent of the metrics used to reduce dimensionality. Hence, from now on we only focus on the PCA results.

### A general mechanism for rhythmic tapping

To dissociate the codification of the passage of time from the parameters that determine the four contexts of the ST, we computed their independent subspaces and the corresponding partial and mixed variance (see Methods). Time encoding explains 38.6% of the total variance, forming circular regenerating loops for each produced interval (Figure 2F, white trajectories). Interestingly, a strong clustering of tapping times was observed when projected in this canonical rhythmic clock (Figure 2F green dots; mean resultant 0.99 circular sd .05; Rayleigh’s test, p < 0.001), emphasizing the notion of an attractor state for triggering the internal beat.

Duration explained 2.7% of the total variance, with subspaces for short and long intervals that were slightly separated (Figure 2F cyan and cinnamon diamonds), due to their increase in amplitude for longer target durations. Modality explained 40.8% of the variance, with very distinct subspaces for visual and auditory conditions since the modality displaced the neural trajectories to a different region of neural state space (Figure 2F red and blue asterisks). Importantly, the mixed variance between time encoding, duration, and modality was 17.9%, indicating that the codification of time shared neural resources across durations and modalities. These findings validate the hypothesis of a strong rhythmic timing machinery in MPC with cyclical neural trajectories that converge in an separatrix at the tapping time while counting the relative passage of time on each regenerating loop, independently of the rget duration, modality, or serial order. Nevertheless, the context also imprints specific signatures in the neural trajectories, as revealed next.

### Kinematics of neural trajectories and modality effects

The kinematics of neural trajectories was characterized using the amplitude, angle, and relative position (Figure 2A-C) between an arbitrary point in state space (fuchsia point Figure 2C) and the neural trajectories across the four ST contexts shown in Figure 2A. We plotted these parameters as a function of the phase (relative timing) of each of the four produced intervals in the sequence (Figure 3).

**Figure 3.**
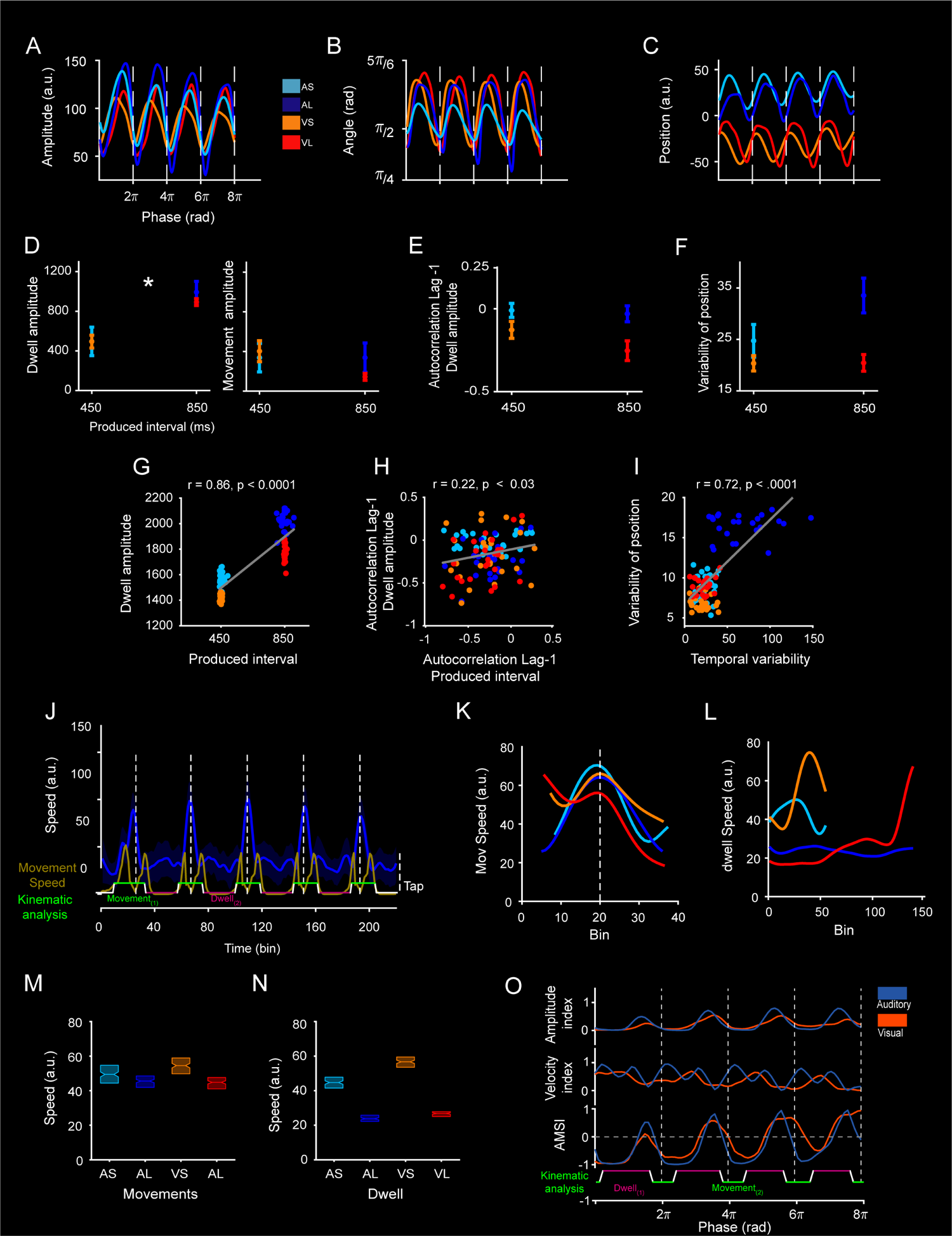
Kinematic of neural trajectories. (A) Amplitude of trajectories computed as the Euclidean distance between an anchor point (fuchsia dot in Figure 2C) and each neural state of the trajectory (Figure 2C) across the four ST conditions. Inset: boxes indicate the ST condition. Dashed lines represent the tap phase. (B) Angle computed from the dot product between the anchor point in A and the neural state of the trajectory. (C) Position computed as the signed difference between an anchor point (fuchsia dot in Figure 2D) and neural trajectories for the four task conditions. (D) Dwell and Movement time amplitude (± 2xSEM) computed as area under curve from A as a function of target interval. The ANOVA showed significant main effects of duration, F(1,192) = 137.42, p < .0001, modality, F(1,192) = 741.58, p < .0001; epoch, F(1,192) = 128.77, p < .0001; as well as significant effects on duration x epoch interaction, F(1,192) = 13585, p < .0001 and modality x epoch interaction, F(1,192) = 1261.6, p < .0001). (E) Lag 1 autocorrelation of the amplitude of the neural trajectories during the Dwell time as a function of target duration. The ANOVA showed significant main effects of duration, F(1,192) = 4.83, p < .02; modality, F(1,192) = 92.8, p < .0001 but not statistical significance on duration x modality interaction, F(1,192) = 1.86, p = .17. (F) Variability of the position (SD within and across trials) from C as a function of target interval (± 5xSEM). The ANOVA showed significant main effects of duration, F(1,192) = 46, p < .0001; modality, F(1,192) = 8.95, p < .0003; as well as significant effects on duration x modality interaction, F(1,192) = 20, p < .0001. (G) Significant correlation between the produced interval and dwell amplitude (r =.88, p < .0001) for recording session 2 of Monkey 2. (H) Significant correlation between the autocorrelation Lag-1 of the produced interval vs autocorrelation Lag-1 of the dwell amplitude (r = .2, p < .04) for the recording session in G. (I) Significant correlation between the temporal variability of the produced intervals and the variability of the trajectory position (r = .72, p < .0001) for the recording session in G. (J) Temporal profile of the speed of the neural trajectory for the four produced intervals of the 850 ms auditory condition in dark blue. Note the large peaks in speed at the tapping times (white vertical dotted lines). The mean speed profile of the hand movement is also showed in the dark yellow trace. At the bottom the subdivision of dwell (pink) and movement (green) periods based on the hand speed are depicted. (K) Speed of neural trajectories during the movement epoch as a function of time across the four ST conditions. (L) Speed of neural trajectories during dwell epoch across time for the four conditions. ANOVA results are described on the main text. (M-N) Box plot (median and interquartile values) for the speed of the neural trajectories between the movement and dwell times, respectively, for each instructed interval and metronome modality. (O) AMSI as a function of trial time for the auditory (blue) and visual (orange) conditions, The dwell and movement periods are depicted at the bottom for the four produced intervals using conventions in J.

Notably, the amplitude, angle, and relative position showed cyclical patterns that were repeated across all serial order elements of the ST sequence (Figure 3A-C). It is important to note that similar kinematic properties were obtained when the arbitrary point was located within a contiguous state space manifold (Figure S2).

The amplitude of neural trajectories was significantly larger for long than short target durations in both the auditory and visual conditions as shown in Figure 2A. In fact, when we computed the area under the amplitude curve during the dwell and movement time periods, we found a significant duration dependent increase in amplitude only in the former (Figure 3D). It is important to emphasize that the individual session analysis showed a high correlation (r > 0.5; p < .0001) between the behavioral dwell times and the amplitude of the trajectories during the dwell in the fourteen sessions of Monkey 2 where we characterized the tapping kinematics with video analysis (Figure 3G; Table 1). Furthermore, individual session analysis revealed that the lag 1 autocorrelation of the amplitude of the trajectories during the dwell was negative (Figure 3E), especially for the condition of 850 ms with a visual metronome, also showing a large correlation in 12 of the fourteen analyzed sessions (Figure 3H) with the autocorrelation of the tapping behavior across the four conditions (r > 0.2; p < 0.0001; Table 1). These findings sustain the hypothesis that the amplitude on the state space trajectories in MPC during the dwell is part of the mechanism that controls the pause between movements to define the tempo in the ST. They also support the notion that the error correction for tap synchronization depends on the adjustments in the amplitude of the trajectories during the dwell of consecutively produced intervals.

**Table 1.** Individual session analysis where we computed the Pearson correlation coefficient between the behavioral (top labels) and kinematic parameters of the neural trajectories (bottom labels) in the fourteen sessions of Monkey 2 with videos on ST behavior. Orange shading is associated with the sessions that had the following three significant correlations: dwell time vs dwell amplitude, temporal variability vs variability of position, and Lag-1 autocorrelation of produced interval vs Lag-1 autocorrelation of dwell amplitude. Sessions in blue shading showed at least one or two significant correlations

| #Session | DWELL TIME |  |  |  | TEMPORAL VARIABILITY |  | AUTOCORRELATION LAG-1 PI |  |  |  | #Cells |
| --- | --- | --- | --- | --- | --- | --- | --- | --- | --- | --- | --- |
|  | Dwell amplitude |  | Dwell speed |  | Variability of position |  | Autocorrelation Lag-1 Dwell amplitude |  | Autocorrelation Lag-1 Dwell speed |  |  |
|  | r | p | r | p | r | p | r | p | r | p |  |
| 2 | 0.59 | 0.0001 | −0.76 | 0.0001 | 0.72 | 0.0001 | 0.23 | 0.02 | 0.26 | 0.01 | 42 |
| 3 | 0.9 | 0.0001 | −0.54 | 0.0001 | 0.18 | 0.0001 | 0.21 | .03 | 0.21 | 0.04 | 28 |
| 6 | 0.55 | 0.0001 | −0.62 | 0.0001 | 0.42 | 0.0001 | 0.2 | 0.04 | 0 | 0.97 | 97 |
| 8 | 0.7 | 0.0001 | −0.41 | 0.0001 | 0.85 | 0.0001 | 0.2 | 0.04 | 0.13 | 0.21 | 76 |
| 9 | 0.86 | 0.0001 | −0.24 | 0.03 | 0.4 | 0.0001 | 0.21 | 0.03 | 0.1 | 0.33 | 84 |
| 11 | 0.85 | 0.0001 | −0.17 | 0.19 | 0.79 | 0.0001 | 0.2 | 0.04 | 0.1 | 0.34 | 75 |
| 14 | 0.92 | 0.0001 | −0.35 | 0.001 | 0.37 | 0.0001 | 0.21 | 0.004 | −0.08 | 0.43 | 21 |
| 1 | 0.95 | 0.0001 | −0.95 | 0.0001 | 0.49 | 0.0001 | −0.03 | 0.78 | 0.05 | 0.6 | 92 |
| 4 | 0.95 | 0.0001 | −0.81 | 0.0001 | 0.49 | 0.0001 | −0.31 | 0.0001 | −0.01 | 0.93 | 97 |
| 6 | 0.93 | 0.0001 | −0.79 | 0.0001 | 0.4 | 0.0001 | −0.17 | 0.09 | 0.08 | 0.44 | 91 |
| 7 | 0.94 | 0.0001 | −0.7 | 0.0001 | 0.35 | 0.0001 | −0.35 | 0.0001 | 0.12 | 0.22 | 94 |
| 10 | 0.98 | 0.0001 | −0.38 | 0.01 | 0.59 | 0.0001 | −0.31 | 0.0001 | −0.06 | 0.54 | 81 |
| 12 | 0.99 | 0.0001 | −0.28 | 0.05 | 0.58 | 0.0001 | 0.09 | 0.37 | −0.04 | 0.68 | 64 |
| 13 | 0.98 | 0.0001 | −0.61 | 0.0001 | 0.62 | 0.0001 | −0.01 | 0.9 | −0.01 | 0.91 | 22 |
| Significant sessions | 14 |  | 13 |  | 13 |  | 7 |  | 1 |  |  |

As expected, the angle acquired minimum values at the tapping times across conditions (Figure 3B), validating the existence of an attractor state that triggers the internal pulse signal when the trajectories reached a specifical angular value (mean resultant 0.99 circular sd 0.11; Rayleigh’s test, p < 0.001).

This experiment had a block design, where each of the four conditions (short, long, auditory, and visual) were recorded for 25 trials before switching to the next, in random order. Hence, the animals knew in which context they were performing ST and this knowledge could act as a tonic external input. In fact, the modality information seems to displace the neural trajectories in different subspaces without greatly altering their cyclical organization, duration dependent changes in amplitude, nor the tap attractor state behavior. This displacement was well captured by the relative position of the trajectories, which showed isomorphic changes within durations of the same modality, but large and significant differences between the auditory and visual conditions (Figure 3C).

Therefore, a modality dependent tonic external input could diverge the cyclic neural trajectories to different subspaces, and the interaction between the amodal rhythmic clock with this external input could shape the differences in temporal precision between the auditory and visual metronomes. Indeed, the trial-by-trial variability of the relative position was highly correlated (Figure 3I) with the temporal variability of the monkeys (r = > 0.35, p < .0001) in 13 of the 14 analyzed sessions of Monkey 2, suggesting that the increase in slope of the scalar property for the auditory metronomes depended on the variability of the neural trajectories within the auditory subspace. Accordingly, the preference for visual metronomes in monkeys and the small associated temporal variability depended on the restrained variability of the neural trajectories within the visual subspace.

To further scrutinize the role of temporal scaling in time encoding, we computed the speed of the neural trajectories across the four ST conditions. Figure 2J shows the complex temporal profile of the speed for the four produced intervals in the long target interval of the auditory condition. Hence, the speed of neural trajectories during the ST does not work as a dial knob to encode a predicted interval, as it has been reported for single interval perception and reproduction (Egger et al., 2019; Wang et al., 2018). In fact, a large repetitive peak of speed occurs at the tapping times, with a relatively steady level between taps. We compared the speed of the neural trajectories between the movement and dwell times. During movement, the speed reached a peak close to the tapping time across durations and modalities (Figure 3K). In contrast, during the dwell the speed showed a larger decrease between the short and long durations (Figure 3L). The corresponding ANOVA showed significant main effects of epoch (F(1,199) = 1368.9, p < 0.0001), duration (F(1,199) = 3137, p < 0.0001), and modality (F(1,199) = 192.5, p < 0.0001), and a large duration x epoch interaction (F(1,199) = 728.8, p < 0.0001). There was a significant negative correlation between the speed and the dwell time of the monkeys across conditions, linking the behavior with temporal scaling (r > .95, p < .0001) in the 14 individual sessions. However, the lag 1 autocorrelation of the speed during the dwell showed significant correlation with the behavioral autocorrelation in only 7 out of the 14 sessions analyzed individually (see Table 1).

The decrease in speed as a function of duration through the dwell supports the hypothesis that time encoding during this critical task period depends on temporal scaling while, as described previously, also depends on changes in the amplitude of the neural trajectories. Nevertheless, the scaling index of the PCs was smaller during the dwell than for the movement time. For the dwell the scaling index was 0.66, 0.4 and 0.73 for the 3 PCs of the auditory condition, and 0.59, 0.64 and 0.54 for the visual condition. For the movement time the scaling index was 0.9, 0.9 and 0.83 for the auditory condition, and 0.72, 0.55 and 0.65 for the visual condition. This apparent contradiction is due to the geometry of the neural trajectories. Even if the speed difference between short and long duration is larger for the dwell (Figure 3L), during this epoch the change in amplitude across target intervals is also larger making the scaling index that focuses on the shape of the trajectories smaller (Figure 3D; Figure S3). Hence, to address this discrepancy we developed an index that determines simultaneously the impact of amplitude and time scaling of the first 3 PCs of the neural trajectories, called the amplitude-modulation-time-scaling index (AMSI). The AMSI is computed as follows:

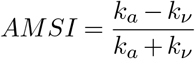

where *k*_*a*_ corresponds to the amplitude index, defined as:

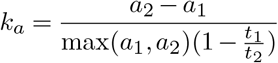

and *k*_*ν*_ correspondes to the velocity index, defined as:

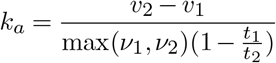

and a, v and t correspond to the amplitude, velocity, and time (target interval) of the neural trajectories for short (1) and long (2) durations (see Figure S4). A logarithmic sigmoid function *σ* was used to adjust *k*_*a*_ = *σ*(*k*_*a*_) and *k*_*ν*_ = *σ*(*k*_*ν*_) to get values within a 0 to 1 range. Thus, AMSI reached a value of −1 when the neural trajectory was fully temporal scaled, while reached a value of one when the neural trajectories were fully amplitude-modulated. The results showed, first, that *k*_*a*_ and *k*_*ν*_ time series showed an oscillatory behavior across the ST, where *k*_*ν*_ attained large values after each tap while *k*_*a*_ showed peak values before each tap (Figure 2O, top-middle). The AMSI was close to −1 at the beginning of the produced interval, increased monotonically within the interval, and reached values close to one just before the next tap (Figure 2O, bottom). Remarkably, AMSI was around zero in the middle of the interval, during dwell time. This behavior was similar between modalities, although the *k*_*ν*_ showed a bimodal behavior within intervals for the auditory condition. These findings indicate that the encoding of the passage of time during the dwell depended on an almost perfect balance between amplitude-modulation and time-scaling across modalities.

A recent algorithm has been developed to test whether the properties of neural trajectories are due to the emergent properties of the neural population or are the result of pooling single neurons. This method compares the neural population responses with surrogate data that preserves simultaneously the temporal correlation of discharge rates, the signal correlations across neurons, and the tuning to the experimental parameters of the task (Elsayed & Cunningham, 2017). We computed the percentage of variance explained by the Duration and Modality projections into the subspaces of Figure 2B and D, respectively. We found that the PC2 of Duration (marginal with p = 0.065) and the PC3 of Modality captured an explained variance that was above the null distribution of surrogate data (Figure S5). These findings suggest that the neural population trajectories provide a concise representation of the rhythmic block during ST, whose collective activity add some emergent information to the representation of rhythmic time, duration, and modality. Consequently, the next step was to focus on the single cell and neural sequence encoding properties of these parameters during the ST.

### Single cell neural encoding

We were interested in testing whether different aspects of the time varying activity of singles cells were related with the key parameters of the task and with the properties of population trajectories. Initially, we carried out a four-way ANOVA using the discharge rate of a cell as dependent variable, and the elapsed time (ET; 20 bins for each produced interval), the duration (Dur; 450 and 850), modality (Mod; Auditory vs visual), and serial order (SO; 1 to 4) of the ST as factors (See Methods). Notably, 86.5% of cells (n = 881) exhibited significant main effects on Elapsed Time and/or the interaction of Elapsed time with the other three factors (see Figure 4A).

**Figure 4.**
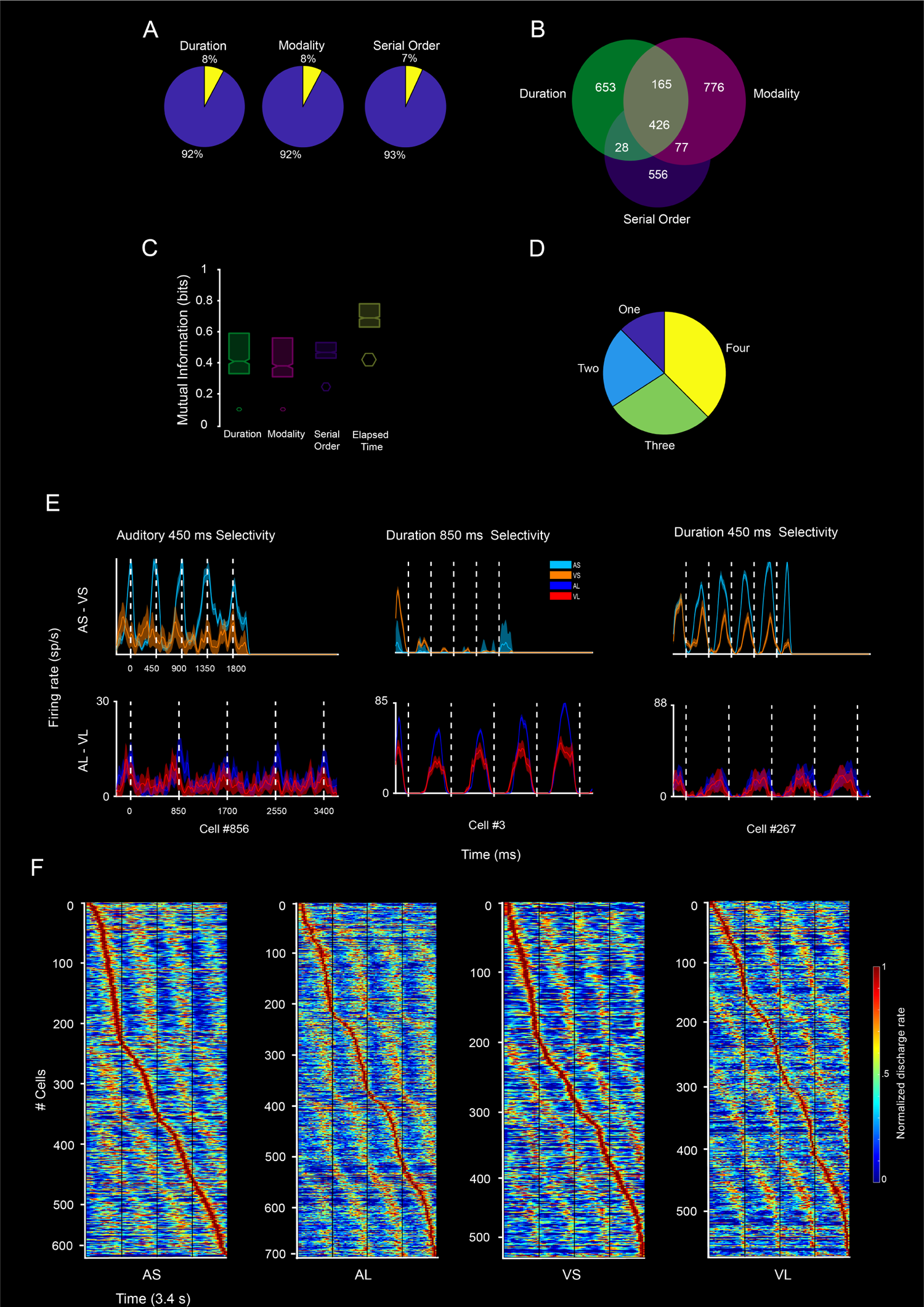
Properties of the neuronal sequences. (A) Pie plots of the proportion of neurons with significant main effects (ANOVA, see main text) on duration, modality and serial order that also had significant (blue) or non-significant effect of elapsed time(yellow). (B) Venn Diagram for number of cells with significant effects for duration (green), modality (magenta), and/or serial order duration (purple). (C) Box plot (median and interquartile values) of the Mutual information for cells with significant permutation test on duration, modality, serial order and elapsed time. The permuted values (median and interquartile values) are showed at the bottom of each parameter. (D) Pie plots of the proportion of cells with significant MI for one to four task parameters. (E) Prototypic examples of neurons with different selective profiles to Modality (left) and Duration (middle and right). (F) Average normalized firing rate of cells (y-axis) with a significant MI on at least one of the ST parameters displayed as a function of trial time for the fours task conditions (AS, AL, VS, VL). The four vertical black lines represent the tapping times. The cells were aligned to the bin of peak activity. Note the dynamic activation of cells throughout the four produced intervals in the ST sequence, with smaller flanking peaks occurring at successive serial order elements of the task.

In addition, Figure 4B depicts the Venn Diagram for cells with significant effects for duration, modality, and/or serial order. Since many neurons showed response modulations for multiple parameters (n = 967), we tested whether MPC neurons showed mixed selectivity. Based on the ANOVA, we divided the neurons into three categories: (i) neurons with classical selectivity (CS; namely no mixed selectivity), which exhibited only a main effect on one of four factors; (ii) neurons with linear mixed-selectivity (LMS), which exhibited main effects between at least two factors but nonsignificant interactions between them; and (iii) neurons with nonlinear mixed selectivity (NMS) which showed significant interaction terms.

We found only 7.3% of CS (n = 75: 48 ET, 9 Dur, 13 Mod, 5 SO) and 5.5% of LMS (n = 56), while 82% of cells were classified as NMS (n = 836). Next, we used mutual information on the binned activity (20 bins for each produced interval) to determine the strength with which the cells represented Elapsed Time, Duration, Modality and Serial Order on the cells with significant effects on the ANOVA. The 94.5% of cells showed significant MI (n = 914/967; permutation test, p < 0.01) for at least one bin and one parameter, with many cells showing large MI above random (Figure 4C) for multiple parameters (Figure 4D). Prototypic examples of neurons with different selective profiles to Modality and Duration are shown in Figure 4E. In general, these findings reveal a large and strong mixed representation for the passage of time with the other task parameters in MPC.

### Sequential patterns of neural activation

Figure 4F shows the average normalized firing rate of the cell population with a significant MI for at least one ST parameter aligned to the bin of peak. Across the four task conditions, there was a dynamic activation of cells throughout the four produced intervals in a sequence, with smaller flanking peaks occurring at successive serial order elements of the task. These cells were recruited in rapid succession producing a progressive neural pattern of activation that flexibly fills each beat duration depending on the tapping tempo, providing a relative representation of how far an interval has evolved. To investigate which properties of the evolving patterns of activation were associated with the internal pulse-like representation of time, as well as with the Duration, Modality and Serial Order, we determined the onset and extent of the activation periods for each cell using the Poisson-train analysis (see Methods). The activation periods of the cells were sorted by their time of peak activity with respect of the previous tap, producing a moving bump for each produced interval and defining four regenerating loops of activation patterns (Figure 5A).

**Figure 5.**
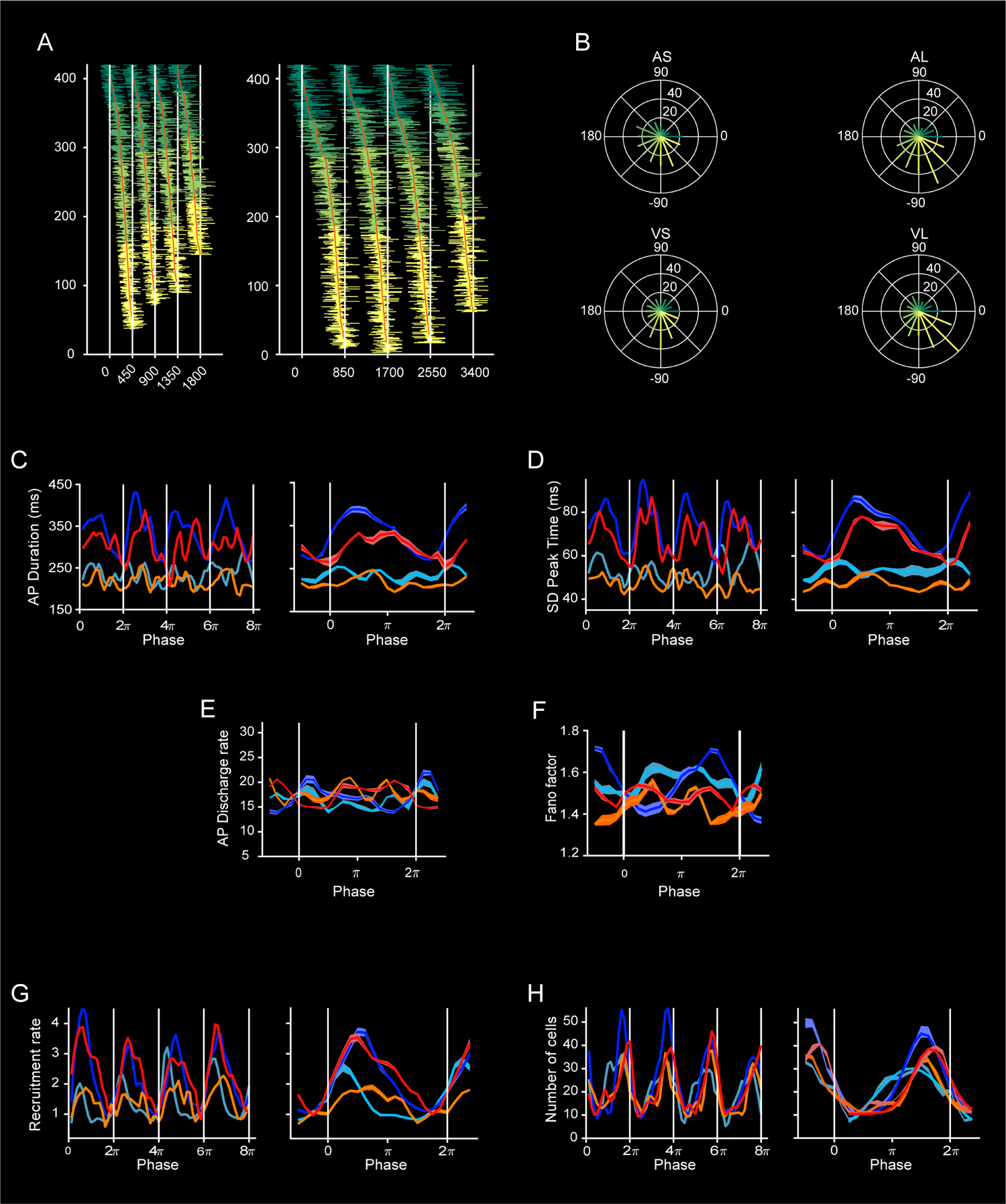
Kinematic of neural trajectories. (A) Amplitude of trajectories computed as the Euclidean distance between an anchor point (fuchsia dot in Figure 2C) and each neural state of the trajectory (Figure 2C) across the four ST conditions. Inset: boxes indicate the ST condition. Dashed lines represent the tap phase. (B) Angle computed from the dot product between the anchor point in A and the neural state of the trajectory. (C) Position computed as the signed difference between an anchor point (fuchsia dot in Figure 2D) and neural trajectories for the four task conditions. (G) Dwell and Movement time amplitude (± 2xSEM) computed as area under curve from A as a function of target interval. The ANOVA showed significant main effects of duration, F(1,192) = 137.42, p < .0001, modality, F(1,192) = 741.58, p < .0001; epoch, F(1,192) = 128.77, p < .0001; as well as significant effects on duration x epoch interaction, F(1,192) = 13585, p < .0001 and modality x epoch interaction, F(1,192) = 1261.6, p < .0001). (E) Lag 1 autocorrelation of the amplitude of the neural trajectories during the Dwell time as a function of target duration. The ANOVA showed significant main effects of duration, F(1,192) = 4.83, p < .02; modality, F(1,192) = 92.8, p < .0001 but not statistical significance on duration x modality interaction, F(1,192) = 1.86, p = .17. (F) Variability of the position (SD within and across trials) from C as a function of target interval (± 5xSEM). The ANOVA showed significant main effects of duration, F(1,192) = 46, p < .0001; modality, F(1,192) = 8.95, p < .0003; as well as significant effects on duration x modality interaction, F(1,192) = 20, p < .0001. (G) Significant correlation between the produced interval and dwell amplitude (r =.88, p < .0001) for recording session 2 of Monkey 2. (H) Significant correlation between the autocorrelation Lag-1 of the produced interval vs autocorrelation Lag-1 of the dwell amplitude (r = .2, p < .04) for the recording session in G. (I) Significant correlation between the temporal variability of the produced intervals and the variability of the trajectory position (r = .72, p < .0001) for the recording session in G. (J) Temporal profile of the speed of the neural trajectory for the four produced intervals of the 850 ms auditory condition in dark blue. Note the large peaks in speed at the tapping times (white vertical dotted lines). The mean speed profile of the hand movement is also showed in the dark yellow trace. At the bottom the subdivision of dwell (pink) and movement (green) periods based on the hand speed are depicted. (K) Speed of neural trajectories during the movement epoch as a function of time across the four ST conditions. (L) Speed of neural trajectories during dwell epoch across time for the four conditions. ANOVA results are described on the main text. (M-N) Box plot (median and interquartile values) for the speed of the neural trajectories between the movement and dwell times, respectively, for each instructed interval and metronome modality. (O) AMSI as a function of trial time for the auditory (blue) and visual (orange) conditions, The dwell and movement periods are depicted at the bottom for the four produced intervals using conventions in J.

The neural sequences were divided into quarters, forming a stereotypic chain of events across conditions, starting after a tap with a group of cells, migrating to other two sets of cells during the timed interval, stopping with the last population quarter before the next tap, and simultaneously resetting to the initial set of cells for the next interval. Next, we used a polar plot representation to determine the number of neurons activated every 22.5 degrees of a circumference representing the duration of the produced interval.

The number of recruited cells significantly changed within an interval (Rayleigh’s test, p < 0.0001, all four conditions), with progressively larger number of cells in the last two quarters of each cycle, peaking before every tap and showing a significant drop in cell numbers for the initial segment of the next regenerating loop (Figure 5B). Therefore, this cyclical migration and resetting of neural sequences across the sequence of isochronous produced intervals seems to be a neural population fingerprint of the rhythmic neural clock in MPC across modalities and tempos.

A crucial question is whether this neural clock used a temporal scaling or absolute encoding strategy. Under the temporal scaling scenario, the activation profile of a neuron is the same between target durations but shrinks for short and elongates for longer tempos, with short and long activation periods, respectively. Thus, under this setting a mean activation period of 200 ms for the target duration of 450 ms should produce an activation period of 377.7 ms for the 850 ms duration. Under the absolute timing strategy, the activation periods are the same across durations, but additional neurons are recruited for longer durations, so that the new neurons are active in the last portion of the interval. Hence, if 300 neurons were conforming a neural sequence in the 450 ms target duration we could expect an extra recruitment of 266 neurons for the 850 ms duration. Based on what we learned from the neural trajectories, it was not surprising to observe a mixed encoding strategy on the progressive neural patterns. The activation periods increased as a function of duration but not with full temporal scaling.

The mean duration for the auditory condition was 232.7±33 ms (mean ± SEM) and 316.7 ± 54 ms for the 450 and 850 target durations, respectively, while for the visual condition were 214.9 ± 23 ms for 450 ms and 300.6±24 ms for 850 ms. Therefore, the scaling indexes for the auditory and visual conditions were 0.72 and 0.74.

On the other hand, the number of neurons recruited in the neural sequences was larger for longer durations, with 335 ± 22 and 398 ± 13 for the short and long intervals in the auditory, and 304 ± 10 and 377 ± 5 for the short and long intervals of the visual condition. The numbers of neurons conforming the neural sequences were around 40 percent less than expected in the long durations if an absolute timing strategy was used. In addition to the shared mixed rhythmic timing strategy, the task modality also imposed changes in the properties of the evolving patterns of neural activity. The number of cells within the circumference of a produced interval showed statistically significant main effects of modality (Chi2(2) = 50.2, p < 0.0001, Harrison-Kanji two-way circular ANOVA) and duration (Chi2(2) = 90.1, p < 0.0001), as well as for the modality x duration interaction (Chi2(1) = 6.7, p = 0.009). Larger number of neurons were recruited for longer intervals, as already noted, and a sharper peak on neurons before each tap was observed for visual rather than auditory metronomes.

### Dynamics of neural sequences

We determined how different properties of the neuronal sequences changed within the progression of each interval in the rhythmic sequence, using the same number of bins for each produced interval to compare task conditions. These parameters are the duration, intertrial standard deviation of the peak time, discharge rate, and Fano Factor of the activation periods, as well as the neural recruitment lapse, and the number of neurons. Importantly, these parameters showed a cyclical variation within each produced interval that was repeated across the four serial order elements of the rhythmic sequence (Figure 5C, F and G, Figure 5C-F).

In fact, no statistically significant effects of serial order were found across all of them (ANOVAs with duration, modality and serial order as factors, p > 0.5). This is a remarkable phenomenon that corroborates that the neural clock is resetting for each interval, with the pulse of the metronome as the unit of measurement, not the absolute time across all the trial.

We carried out ANOVAs on each parameter using Duration, Modality, and the Quarter of the produced interval as factors. We found that the duration of the activation periods was larger for auditory than visual metronomes, and greater for longer tempos, with extended periods of activation in the middle two quarters of the 850 ms intervals (Figure 5C). Accordingly, the standard deviation of the peak time was larger for the 850 ms intervals of both modalities, with lower values around the tapping times (Figure 5D). Importantly, no evidence of an increase in peak response variability as a function of absolute time was obtained, rejecting the possibility of a Weber law scaling of response variability over absolute time (Cao et al., 2022). The discharge rate of the activation periods was slightly larger for the visual than the auditory condition, especially in the last two quarters of the produced intervals (Figure 5E). The Fano Factor, a coefficient of variation in the neural responses during the activation periods, was larger for the auditory than the visual condition on the last two quarters of the produced interval (Figure 5F).

The neural recruitment lapse, which is the time between pairs of consecutively activated cells, showed a larger increase in the first quarter that plateaued until the third quarter and acquired lower values on the last quarter. The magnitude of this cyclical pattern was larger for the longer target interval of both metronome modalities (Figure 5G). Finally, the number of cells showed an initial decrease followed by rebound that was steeper for the visual condition, and a peak in the fourth quarter followed by a sharp decrease at the end of the produced interval. These results confirm a mixed coding strategy for rhythmic timing with time scaling, absolute timing, and modality specific components. Furthermore, the larger recruitment of cells during the first three quarters for longer durations allows the generation of neural sequences that cover all the 850 ms interval, building a relative timing clock that engages cells migrating at different speeds to fill the interval. On the other hand, the sudden increase in number of cells just before the tap is a crucial event that could signal the internal prediction of the metronome’s pulse (Figure 5H). At the level of the neural trajectories, the peak in the number of cells can be linked to both the tap separatrix and the peak in the speed of neural trajectories. These hypotheses are tested in the simulations described below.

Cannon’s model of rhythm tracking (Cannon, 2021) posits a neural process that acts as an estimate of the phase *ϕ* of a cyclically patterned stimulus. The cyclical neural sequences with mixed time coding strategies could be the neural correlate for this phase estimating process. In this model, the expectation of sensory events at each phase is quantified by a function *λ*(*ϕ*) that takes the form of a sum of Gaussian functions with means *ϕ*_*i*_ representing the phases at which events are expected, variances *v*_*i*_ representing the imprecision of predicted times, and scaling factors *λ*_*i*_ encoding the strength of the expectations. The sudden recruitment of cells before the tap could represent a peak in this expectancy function. To attempt to determine whether the cyclic variation in the number of recruited cells is a neural correlate of this predictive point process, we fitted a Gaussian function on the number of cells as a function of time bin. The fittings revealed that, according to this interpretation of cell recruitment, the expected internal beat was close to the end of the interval across conditions (means close to 2*π*), the expectation was less precise for auditory than visual metronomes and larger for longer target intervals, and the strength of the expectations was larger for slower tempos (Figure 5H, left). These properties define a phasic temporal expectation signal that is modulated by tempo and modality.

It is important to mention that the neurons within the neural sequences showed instantaneous activity changes that correspond to different types of ramping patterns that have been reported previously (see Figure S6) (Henke et al., 2021; Knudsen et al., 2014; Merchant et al., 2011; Merchant & Honing, 2014). The neurons of the first quarter of the moving bumps showed the ramping profile of the swinging that encode the beginning of each produced interval. The neurons of the second and third quarters showed an up-and-down profile of activation that is characteristic of the ramps that encoded elapsed time in the width of the ramp in a rhythmic tapping task (Merchant et al., 2011; Merchant & Averbeck, 2017). Finally, the neurons of the last quarter showed ramping activity that is distinctive of cells encoding the time-remaining for an action, reaching a peak of activity at a particular time before the tap across conditions, with larger slopes for shorter durations (Merchant et al., 2004b, 2011).

### Generalizability of neural sequences

We plotted the normalized activity of the 918 neurons that showed significant effects in the encoding across the four task conditions, sorting each condition according to the latency of peak activity for every unit of one task and using this sorting across tasks. These heatmaps were constructed with the cell response profiles segmented in 20 bins and averaged across the four serial order elements of the ST, for each of the four task conditions. Clear gradual patterns of activation were observed of self-sorted moving bumps (diagonal panels Figure 6A), while cross-sorted sequences were quite different, with complex patterns of activity suggesting small generalizability of neural sequences between durations and modalities (off-diagonal panels Figure 6A). To determine quantitatively the change in the dynamics of neural sequences across conditions we adopted a geometric approach that has been used before Remington et al. (2018); Zhou et al. (2022). First, we computed the Euclidean distance matrix between all possible pairs of task conditions using the sorted activity of Figure 6A. Second, we identified the minimum distance value for each bin within each distance matrix and extracted two vectors: one vector keeps the distance values at the minimum (called the minimum distance vector), and the other keeps the bin difference between the diagonal and the minimum values (called the diagonal asymmetry vector). We transformed diagonal asymmetry vector into angular differences, where a difference of 20 bins between the diagonal and the minimum value corresponds to 360 degrees. For the self-sorted neural sequences, the minimum distance and the diagonal asymmetry vectors were constituted by twenty zeros, one for each time bin, since we were comparing a condition with itself (Figure 6B, white dots in task comparison diagonal). In contrast, for cross-sorted moving bumps the minimum values of the distance matrix can fall in different values between the rows and columns and both vectors were different from zero (Figure 6B, white dots in task comparison off-diagonal).

**Figure 6.**
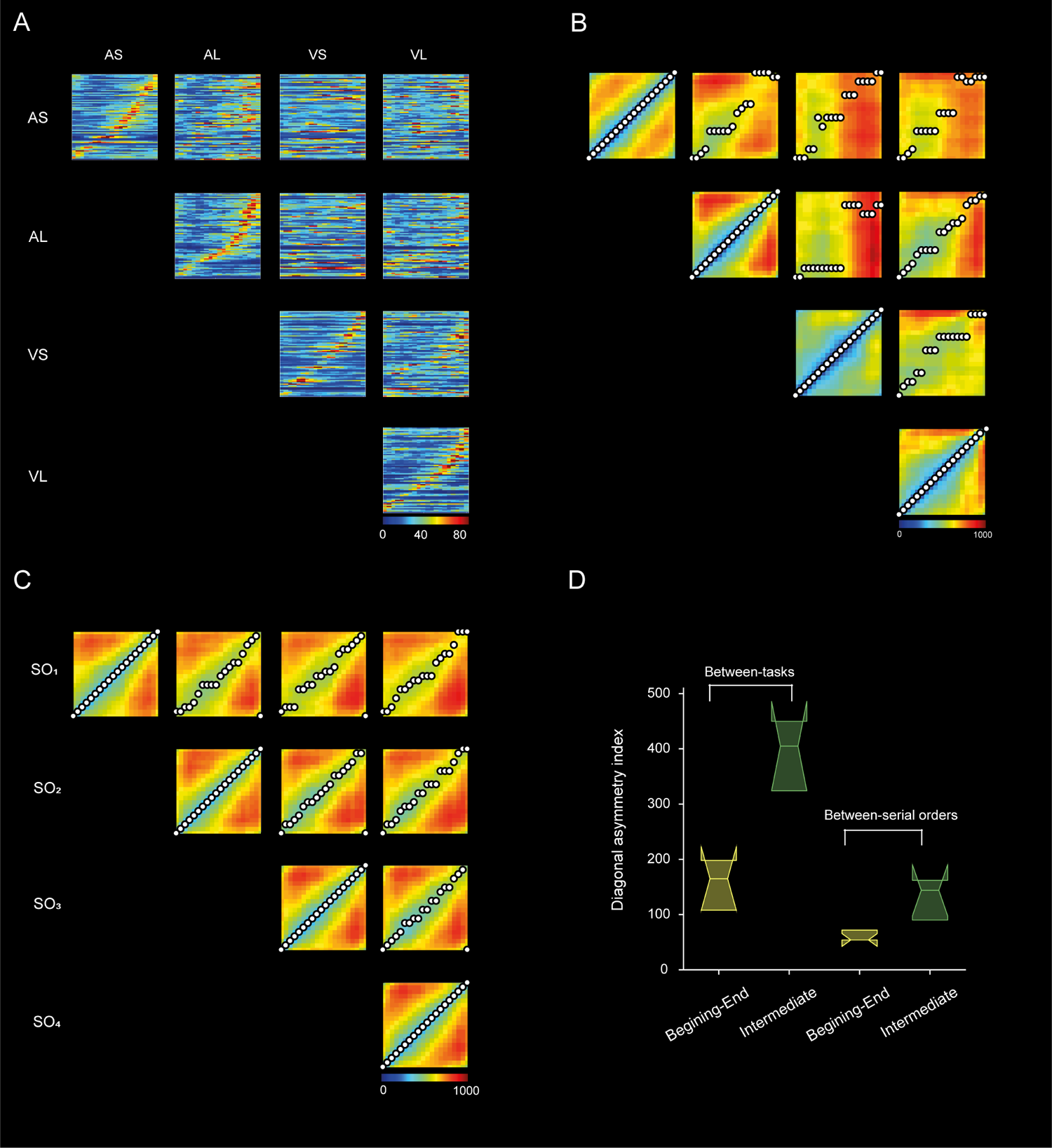
Generalizability of neural sequences. (A) Gradual patterns of activation of self-sorted (diagonal panels) and cross-sorted (non-diagonal panels) moving bumps. Cross-sorted sequences show complex patterns of activity suggesting small generalizability of neural sequences between durations and modalities. Task conditions are depicted at the edges: AS auditory short, AL auditory long, VS visual short, VL visual long. The neural activity is depicted as discharge rate over time with 20 bins for each produced interval. (B) Euclidean distance matrix between all possible pairs of task conditions using the sorted activity of A. For the self-sorted conditions (diagonal panels) the Euclidean distance is zero. The minimum distance per bin is depicted as a white circle. Same conventions as in A. (C) Euclidean distance matrix between serial order elements of the auditory long condition. Same conventions as in B. (D) Diagonal asymmetry index for initial intermediate and final bins using the Euclidean distance across task conditions (left, from panel B) and serial order elements of the auditory large condition (right, from panel C). Similar results were obtained for serial order distances using neural sequences of the other three ST conditions (data not shown).

Third, we defined the distance index as the mean of the minimum distance vector, and the diagonal asymmetry index as the sum of the diagonal asymmetry vector. The former is a measure of overall distance between the neural sequences of a pair of task conditions, whereas the latter is a cumulative measure of how congruent over time are the two neural sequences within each produced interval of the ST. The results showed that both the distance and the diagonal asymmetry indexes were larger between than within modalities and larger between than within durations (Figure 6B, see Figure S7), suggesting strong effects of both task parameters in the cell configuration and dynamics of the neural sequences. Since the white dots in the distance matrices of Figure 6B showed larger diagonal asymmetry for intermediate than for the initial and final bins, we also computed this index for the two periods. Indeed, the diagonal asymmetry index for intermediate bins was quite larger and statistically different from the corresponding index for initial-final bins (Figure 6C; F(1,19) = 5.02, p < 0.048; Watson-Williams circular test), indicating that the neurons that were active within the interval were more context dependent that the cells after or before the taps. To further test this notion, we first computed the correlation of the activation profile between pairs of cells sorted as in Figure 6A. Interestingly, the values for the correlation matrix of self-sorted cells for auditory short condition were not only large around the diagonal but also for pairs of cells that were active at the beginning and end of the neural sequence (Figure S7A). Then, we computed the probability that pairs of cells showed large correlation values (r > 0.5) in their response profile dividing the complete neural sequence into quarters (Figure S7B). We found large probability values for the last two quarters of the moving bump across all the possible comparisons of task conditions, supporting the existence of a neural population that is similarly active close the next tap across durations and modalities (Figure S7B). In fact, we found 124 neurons with similar activation profiles across the four conditions and the four serial order elements of the ST, which were active mainly at the end of the produced interval (Figure S7C).

We also computed the distance and the diagonal asymmetry indexes for neural sequences sorted by the serial order elements within each ST condition. Figure 6C depicts the Euclidean distance matrices for neural sequences of the long auditory condition sorted by serial order elements, with the minimum distance values as white dots. There is a systematic degradation in the neural sequence organization as the difference in serial order increases, accompanied by an increase in both indexes. In addition, the distance and diagonal asymmetry indexes were smaller for serial order comparison than task conditions (see Figure 6D vs Figure S7D). We carried out a two-factor circular ANOVA using the diagonal asymmetry index as dependent variable, and the bin-epoch (intermediate vs. initial-final bins) and the distance configuration (task vs the serial order) as factors (see Figure 6C). The results showed no significant main effects but a significant bin-epoch x configuration interaction (Figure 6C; 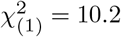, p < 0.0014; Harrison and Kanji two-way ANOVA for circular data). In addition, the distance index was significantly larger for task condition than serial order, but similar between bin-epoch (ANOVA, distance configuration F(1,20) = 0.22, p = .88; bin-epoch F(1,20) = 13, p < .001; distance configuration x bin-epoch interaction F(1,20) = 0.35, p = .55). These findings suggest that there was a stable and quite repetitive pattern of activation in the moving bumps across the sequential structure of the task, which is different from the strong reconfiguration of the neural sequences between modalities and durations, which specially occurred within the produced interval.

Overall, these findings support the theory that neural sequences during the ST form a congruent wave of neural activation that regenerates for each rhythmically produced interval, engaging similar evolving patterns of activity across consecutive intervals. Furthermore, the moving bumps recapitulate many properties of the neural trajectories, including: (1) cyclical patterns that are renewed across the sequential structure of ST; (2) generating relative instead of absolute time representation; (3) the activation of larger groups of cells close to the taps signaling an internal pulse and then resetting the moving bump, most of these cells are active across durations, modalities and serial order elements; (4) the combination of an increase in the number of engaged neurons, larger recruitment lapses between neurons, and a rise in the duration of their activation periods as a neural foundation for the mixed representation of tempo in the amplitude modulation and temporal scaling of the neural trajectories; (5) the small generalizability between modalities and durations that suggest different inputs to the MPC for auditory and visual metronomes.

### Relations between neural trajectories and neural sequences

We simulated evolving patterns of population responses with specific profiles of activation and evaluated their conversion to state space by projecting the activity in three PC dimensions. This allowed testing different hypothesis regarding the effect of the key properties of the moving bumps just mentions above on the AMSI and separatrix behavior for taps on the neural trajectories.

Initially, we simulated neural sequences with an increase in the number of neurons for the longer duration, but similar duration and magnitude of response between the two target intervals and no resetting in the moving bumps for taps. The resulting moving bumps corresponded to an absolute neural clock that encodes elapsed time from the beginning to the end of the trial on the final position of the neural trajectories (Merchant & Pérez, 2020; Zhou et al., 2022). There is no rotatory component since there is no resetting in the neural sequences, and the corresponding AMSI showed values above 0 indicating changes in amplitude (right column Figure 7A) but not in temporal scaling. Next, we simulated absolute but resetting moving bumps for two produced intervals, with cyclical trajectories that showed no temporal scaling just changes in amplitude (AMSI with values around 1) (Figure 7B). Conversely, relative resetting neural sequences, with the same number of neurons and a scaled increase in the response duration, produced cyclical neural trajectories with AMSI around −1, indicating pure temporal scaling and no changes in amplitude (Figure 7C). If instead of changing the duration we change the magnitude of the response on the longer interval in the previous simulation, the AMSI showed values around 1 indicating amplitude modulation (Figure 7D). Hence, changes in duration of the activation period produces temporal scaling, while changes in the number of cells and response magnitude produced changes in the amplitude of neural trajectories, as seen in the empirical data (Figures 3 and 5).

**Figure 7.**
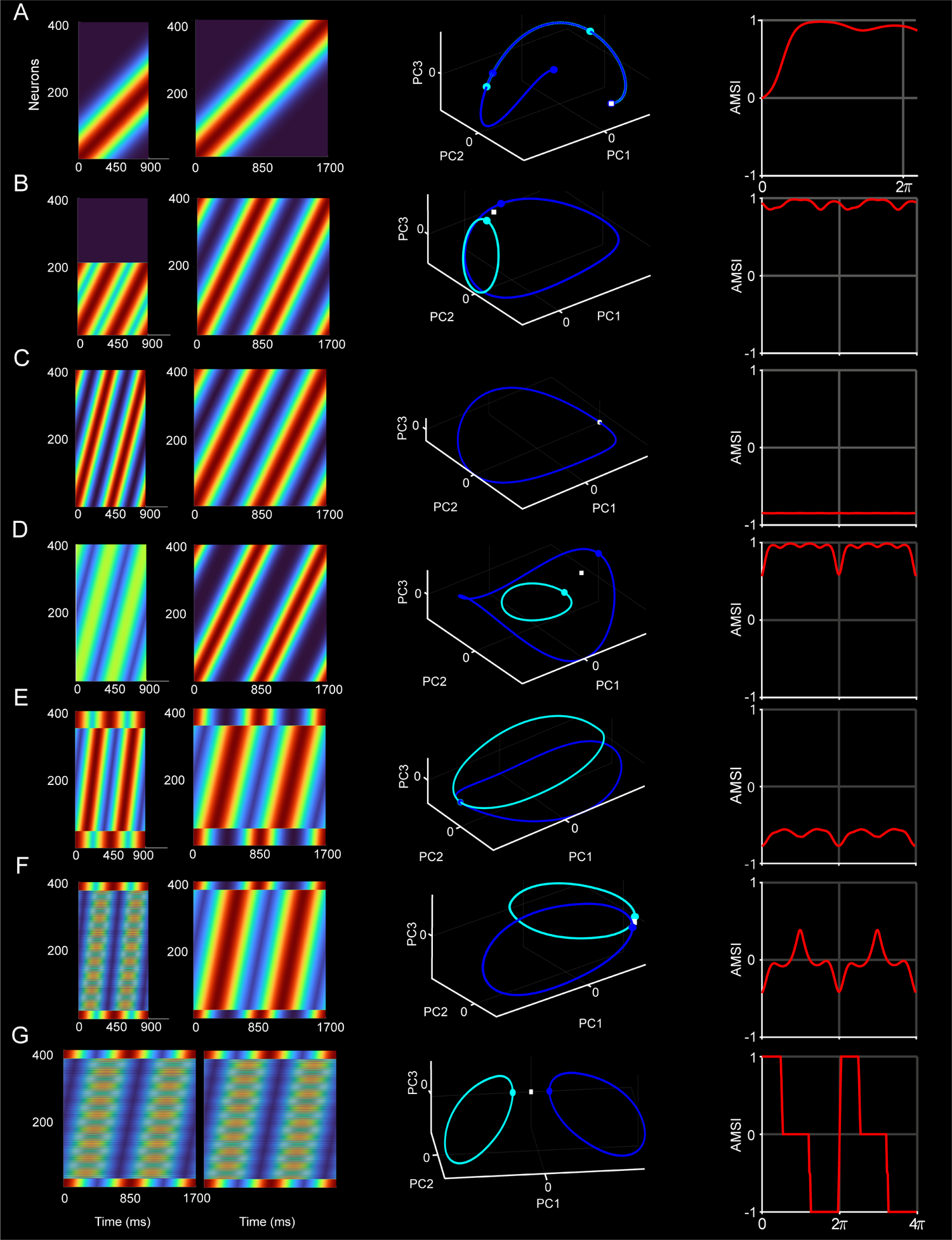
Neural sequence simulations and their resulting neural trajectories. (A-G) First two left columns: Heatmaps of the normalized activation profiles for short and long intervals. Middle: neural trajectories for the short (cyan) and long (dark blue) conditions. Right: AMSI as a function of time or tapping phase. (A) Neural sequences for a single interval with the double number of neurons for the long interval, but the same duration and magnitude on the responses. (B) Same as in A but for two resetting neural sequences simulating two chains of activation for two rhythmic intervals. (C) As in B but with the same number of neurons, a temporally scaled activation period, and same response magnitude. (D) As in C with the same number of neurons, same response duration and larger response amplitude for the long interval. (E) Neural sequences sharing neurons at the beginning and end of each produced interval and a temporally scaled response. (F) Same as in E, with a temporally scaled activation period and larger number neurons for the long interval. (G) Neural sequences with the same duration and shared neurons at the beginning and end of the two produced intervals but with partially overlapping cells within intervals.

We found that sharing neurons at the beginning and end of each produced interval across target intervals, emulating the observed moving bump organization (Figure 6 and S8), induced a convergence of the neural trajectories at tap times and, hence, a separatrix behavior (Figure 7E). Indeed, the distance of the trajectories at tapping times is zero or very small when simulations include a shared population of cells at the beginning and end on the interval (Figure S8). These findings confirm the notion that the neural internal pulse representation as the tap separatrix depends on the activation of a group of neurons whose activity flanks the interval produced interval, responding similarly across durations, modalities, and serial order elements. Adding to this simulation an increment in duration of the activation periods, as well as larger number of neurons in the intermediate epoch of the interval for longer durations, produced neural trajectories that were similar to the original population dynamics (Figure 7E and 7F). These neural trajectories were circular, they converge at tapping times, and showed cyclical variations of AMSI with values close to zero in the middle of the produced interval, indicating a mixture of temporal scaling and amplitude modulation for time encoding. This combined simulation supports the deep relationship between the properties of the original neural trajectories and their corresponding neural sequences. Finally, simulating neural sequences with the same set of neurons (Figure 7E), partially overlapping (Figure 7F), or duration-selective populations (Figure. 7G), produced an angular difference in the circular trajectories that was between zero (same neural populations) and 90 degrees (completely different sets of cells). These results suggest that the angular difference in the subspaces of the auditory and visual conditions was largely due to the partially overlapping bimodal cells of our database.

## Discussion

The parametric account of the rhythmic behavior of the animals during the ST, combined with both the geometric and kinematic assessment of neural trajectories, and the detailed analysis of the properties of the neural sequences revealed a series of fundamental principles governing the neural rhythmic clock in MPC. On one side, there is an amodal representation of beat based timing that includes at least three components. First, the trajectories converge in similar state space at tapping times while the moving bumps restart at this point, resetting the beat-based clock. The tap separatrix and the neural resetting is a neural correlate of the internal pulse that coincides with the tapping times, providing a phasic representation of a cyclic time event, as well as a continuous and relative lecture of how much time has passed within each produced interval. Second, the tempo of the tap synchronization depends on the dwell between stereotyped movements. This dwell is encoded by a combination of amplitude modulation and temporal scaling in the neural trajectories, which at the moving bump level correlate with a mixture of an increase in the number of engaged neurons, larger recruitment lapses between neurons, and a rise in the duration of their activation periods. Third, the mechanism for error correction that maintains tap synchronization with the metronome depends on the within trial changes in amplitude of the trajectories during the dwell period. Thus, a longer produced interval with a large amplitude in the neural trajectory tends to be followed by a shorter interval with small amplitude, while a shorter interval with small amplitude tends to be followed by a longer produced duration and the corresponding large amplitude in the neural trajectory. Conversely, the modality of the metronome produced profound changes in the monkeys’ rhythmic behavior, with timing that is more precise, more accurate, and with tap synchronization that showed larger error correction when using visual rather than auditory isochronous stimuli. Accordingly, the modality imprints specific signatures in the neural trajectories, with a large displacement in state space without greatly altering their cyclical organization, duration dependent changes in amplitude and temporal scaling, nor the tap separatrix behavior. These findings suggest the existence of a modality dependent tonic external input that produces a divergence in the cyclic neural trajectories to different subspaces. We also found modality selective cells and a lack of generalizability of the neural sequences between modalities, results that support the notion of a differential input for visual and auditory signals in MPC.

The notion of an internal representation of pulse implies the existence of a phasic signal that is a cognitive construct of the regular temporal expectations based on the properties of the input sequence (Cannon, 2021; Jones & Mcauley, 2005). In the human literature a framework that has been very popular is the Dynamic Attending Theory, which states that rhythmic temporal expectancy depends on pulses generated by coupled oscillators (Large & Jones, 1999; Large & Palmer, 2002). Models using coupled oscillators can explain beat perception from a large variety of rhythms with a complex metric structure (Large et al., 2018). Nevertheless, this approach is built on the strong assumption that the brain works as a generator of long-lasting oscillations that can be coupled in time to represent the pulse. Instead, cortical and subcortical oscillatory activity usually occurs in short burst (200 to 1000 ms) and depend on local inhibitory mechanisms that generates alternating temporal windows of enhanced and decreased excitability (Merchant et al., 2012; Cadena-Valencia et al., 2018). Importantly, this inhibition-based mechanisms produces natural temporal windows to group neuronal activity into cell populations and neural (Buzsáki, 2006; Buzsáki & Watson, 2012). Here, we offer a neurophysiological account for the interval representation of pulse, where the neural population generates a cyclic pattern that provided a continuous and relative lecture of how much time has passes within each produced interval in the rhythmic sequence. This relative time encoding is observed in both the neural trajectories and the neural sequences. In addition, the line separatrix in the neural trajectories at tapping times is a robust neural correlate for internal beat. An ideal reader in state space can naturally generate a tick pulse every time the neural trajectories reach the separatrix. In addition, the sudden increase in number of cells just before the tap is a crucial event that could also signal the internal prediction of the metronome’s pulse (Figure 5H). A recent study modeled internal pulse prediction as a probabilistic point process where an observer is continuously inferring the phase of the pulse based on phasic temporal expectations template that was modelled as the sum of Gaussians (Cannon, 2021). Crucially, the large recruitment of neurons before the tap can be seen as the temporal expectation signal, where the variability of the predicted times was larger for auditory than visual metronomes and larger for longer target intervals, and the strength of the expectations was larger for longer tempos (Figure 5H). These properties correlated with the larger temporal variability for the auditory condition and high accuracy in temporal production for longer intervals (Figure 1). Furthermore, a large proportion of these cells were active before every tap across serial orders and modalities, which coincides with the large generalization of the neural sequences before the tap (Figure 6C). Thus, the line separatrix that is orthogonal to the circular loops in the neural trajectories (Figure 2C) may depend on this last type of cells.

We have shown that independently of the modality monkeys produce rhythmic intervals in synchrony with a metronome by controlling the dwell time between stereotyped tapping movements (Donnet et al., 2014; Gámez et al., 2018). The temporal control of dwell duration depends on a dynamic combination of temporal scaling and amplitude modulation in the neural trajectories. Temporal scaling as a timing mechanism has been reported in MPC (Merchant et al., 2011; Merchant & Averbeck, 2017; Remington et al., 2018; Wang et al., 2018), prefrontal (Henke et al., 2021; Tiganj et al., 2017; Xu et al., 2014); parietal cortex (Jazayeri & Shadlen, 2010; Merchant et al., 2004b), as well as in the basal ganglia (Bakhurin et al., 2017; Emmons et al., 2017; Gouvêa et al.; Mello et al., 2015; Wang et al., 2018). The main concept behind scaling is that the same population of cells represent time to an event (normally a movement) by shrinking or expanding the neural response profile for short or long intervals, respectively. This change in the pattern of activation at the single cell level, produces a decrease in the speed but no differences in geometry on the neural trajectories as a function of the length of the quantified time (Bi & Zhou, 2020; Merchant & Pérez, 2020; Wang et al., 2018). Therefore, the activation speed of the neural population clock is set at a constant level in order to predict and produce a specific duration in single interval tasks (Sohn et al., 2019). In contrast, the speed of the neural trajectories during rhythmic tapping is not constant, reaching a large peak at the tapping time across tempi but showing a decrease for long durations during the dwell (Figure 3E-F). Furthermore, the temporal scaling of the neural trajectories during the dwell was accompanied by an amplitude modulation in their circular changes in state space (Gámez et al., 2019). Importantly, both the amplitude and the speed change are robustly correlated with the intervals produced by the monkeys. Hence, the mixed coding strategy where neural populations combine temporal scaling and amplitude modulation seem to be an especial signature of rhythmic timing. At the neural sequence level, the changes in the duration of the cells’ activation periods were related with temporal scaling, while the number of cells in the sequences and the increase in their recruitment lapse were linked with the amplitude modulation of the neural trajectories. The changing numbers of cells recruited in the moving bumps is an indication of interval tuning. In fact, the mixed selectivity of single cell responses across task parameters was accompanied by a large number of neurons with selective responses for short and long intervals (Figure 4E). Interval tuning during single interval and beat based timing has been reported in medial premotor areas (Merchant et al., 2013b; Mita et al., 2009), prefrontal cortex (Henke et al., 2021)(Henke et al., 2021), the putamen (Bartolo et al., 2014; Bartolo & Merchant, 2015), the caudate (Kameda et al., 2019; Kunimatsu et al., 2018) and the cerebellum (Ohmae et al., 2017; ichi Okada et al., 2022). In addition, a chronomap in the medial premotor cortex has been described in humans using functional imaging, where interval specific circuits show a topography with short preferred intervals in the anterior and long preferred intervals in the posterior portion of SMA/preSMA (Protopapa et al., 2019). Hence, timing not only depends on one population of cells that contracts of expands their activity patterns depending on a constant speed knob, but also on interval specific neurons that generate distinct timing circuits. Tuning and modularity are mechanisms for division of labor that are widely used in cortical and subcortical circuits to represent sensory, cognitive and motor information (Hubel, 1977; Georgopoulos et al., 2007; Goldman-Rakic et al., 1984; Mountcastle, 1998; Naselaris et al., 2006). Interval tuning can provide large flexibility to encode the passage of time and to predict events across behaviors that require the integration of timing with other task parameters that have a different mapping framework in MPC (Garcia-Saldivar et al., 2021; Yu et al., 2005). Since the width of cell tuning is wide, interval tuned neurons can also show temporal scaling (Crowe et al., 2014; Henke et al., 2021; Merchant et al., 2013a), which can be the substrate of the observed mixed timing encoding that combines amplitude modulation and temporal scaling.

It has been shown in humans that performance to an auditory metronome is more precise and accurate than synchronization to a flashing visual metronome (Chen et al., 2002; Merchant et al., 2008b,a; Repp & Keller, 2004). Conversely, macaques showed a bias towards flashing visual metronomes, with rhythmic timing that was more precise, more accurate, and with larger error correction than with auditory isochronous stimuli (Figure 1). This interspecies difference may depend on the anatomofunctional properties of their audiomotor system, especially in parietal cortex that is an intermediate processing node (Honing & Merchant, 2014; Mendoza & Merchant, 2014; Rilling et al., 2008). Posterior parietal cortex processes multimodal information (Andersen, 1997; Cohen & Andersen, 2002) and that is deeply involved in sensorimotor control (Battaglia-Mayer & Caminiti, 2019). In humans, the posterior auditory cortex is amply connected with parietal cortex, forming the dorsal auditory stream for the localization and timing of sound (Ortiz-Rios et al., 2017; Rauschecker, 2018; Schubotz et al., 2003; Woods et al., 2006). Conversely, in monkeys the homologous posterior medial and lateral belt areas of the auditory cortex (Rozzi et al., 2006) only send restricted connections to area 7a and VIP of the parietal lobe. This limited auditory input contrast with the massive reciprocal link between parietal and visual areas in macaques, constituting the dorsal visual stream for spatial, temporal and motion processing (Battelli et al., 2007; Mishkin et al., 1983). For example, area 7a is strongly connected with visual areas that map the foveal and specially the peripheral visual field (V2, V3, PO), as well as with areas involved in visual motion (MT, MST) (Baizer et al., 1991; Cavada & Goldman-Rakic, 1989b,a). Hence, we suggest that the bias towards visual metronomes in monkeys is rooted on their largely visual parietal region, while the remarkable human abilities for auditory beat perception and entrainment depend on their vast audioparietal link.

A fundamental result of the present study is that the modality of the metronome produced a large displacement of the neural trajectories in state space without considerably altering their internal pulse representation or the rhythmic time keeping mechanism. These findings suggest the existence of a modality dependent tonic external input that produces a divergence in the cyclic neural trajectories to different subspaces during ST. A similar mechanism to encode two timing gain contexts for interval reproduction, based on different levels of a static input while preserving the computational timing mechanism, was recently reported in MPC by Jazayeri’s group (Remington et al., 2018). Posterior parietal areas, especially area 7a, could provide this differential input. Indeed, preSMA receives parietal inputs mainly from area 7a (Luppino et al., 1993). This ample region of the posterior parietal cortex receives differential auditory and visual inputs (see above), and is implicated in sensorimotor coordination during reaching, as well as in timing of single and rhythmic intervals (Merchant et al., 2003, 2004a; Merchant & Georgopoulos, 2006; Ross et al., 2018). Consequently, we postulate that area 7a send partially overlapping auditory and visual inputs to MPC and that the uneven connectivity between the two modalities imposes the tonic divergence of the cyclical neural trajectories in the different auditory and visual subspaces. We emphasize the notion of partially overlapping audiovisual inputs to MPC due to both the observed none-orthogonal subspaces for visual and auditory metronomes and the single cell mixed selectivity across modalities (See simulations in Figure 7G).

Lastly, error correction is a critical component of tapping synchronization and is constituted by a phase correction mechanism, involved in subtle corrections of the relative phase between the metronome and the taps, and by a period correction mechanism that adjusts deviations of the internal neural clock period (Jantzen et al., 2018; Repp, 2000). The latter is captured by autocorrelation structure of adjacent taps where a negative lag 1 implies period correction (Iversen et al., 2015). Consistent with the human auditory bias for and the monkey visual bias for beat based timing, the period correction in humans is larger for auditory than visual metronomes (Comstock et al., 2018), while we observed the contrary on monkeys (Figure 1E-F). EEG experiments in humans have shown that MPC is involved in error correction during natural tap synchronization (Bavassi et al., 2013), and during tapping perturbations (Jantzen et al., 2018). Our individual session results showed that the lag 1 autocorrelation of the amplitude of MPC neural trajectories was negative during the dwell, especially for the condition of 850 ms with a visual metronome. Thus, there was a significant correlation of the lag 1 autocorrelation between the tapping behavior and the amplitude of the neural trajectories during dwell in most of the recording sessions with video kinematic analysis. In contrast, more than half of the sessions did not showed significant correlations of the lag 1 autocorrelation between behavior and the speed of neural trajectories during dwell. These results support the notion that although time encoding during dwell depends on a mixed signal of amplitude modulation and temporal scaling, the mechanism behind error correction mainly relies on changes of amplitude. We suggest that the amplitude neural reader may also be engaged in error correction, imposing opposite changes in the amplitude of consecutive intervals to keep tapping in synchrony with the metronome.

## Material and Methods

### Subjects

All the animal care, housing, and experimental procedures were approved by Ethics in Research Committee of the Universidad Nacional Autónoma de México and conformed to the principles outlined in the Guide for Care and Use of Laboratory Animals (NIH, publication number 85-23, revised 1985). The two monkeys (M01 and M02, Macaca mulatta, one male the other female, 5-7 kg BW) were monitored daily by the researchers and the animal care staff to check their conditions of health and welfare.

### Tasks

Synchronization Task (ST). The ST has been described before (Zarco et al., 2009). Briefly, monkeys were trained to attend to a sequence of brief stimuli with a constant interstimulus interval and push a button in synchrony with the latter six of them (Figure 1A). At the beginning of a trial, participants held a lever and attended to three stimuli after which they started to move, the goal being to produce six taps in synchrony with the six remaining metronome pulses. Trials were separated by a variable intertrial interval (1.2–4 s). The inter-onset intervals (IOIs, 450 or 850 ms) for the visual (red square with a side length of 5 cm, presented for 33 ms) or auditory (white noise with a 33 ms duration) metronomes were presented in blocks of 25 trials. Monkeys received a reward (fruit juice) when the duration of the produced intervals showed an error below the 18% of the instructed interval and all asynchronies between stimuli and taps were less than ±200 ms. The order of the two interval/modality combinations was random across days.

### Neural recordings

We used two semichronic, high-density electrode systems (Mendoza et al., 2016) placed bilaterally in the limit between SMA and preSMA, with 64 recording sites (Buzsaki64-Z64) in the left and 64 in the right hemisphere of the MPC. The probes were connected to a microdrive that allowed the control of the movement of the two electrode systems independently in the dorso-ventral axis. The neural data of 128 channels was acquired, amplified, and digitized using a PZ2 preamplifier (Tucker-Davis Technologies, FL, USA, http://www.tdt.com) at 24,414Hz. The signal was transmitted to a RZ2 base station through fiber optic for on-line processing.

### Spike detection discrimination

We developed a spike sorting pipeline (ABVA) which is based on previous algorithms for massive spike discrimination (Rossant et al., 2016). Briefly, it considers that high-density silicon probes can record the action potential of the same neuron in different recording sites (up to the 8 recording sites for each silicon probe), producing simultaneous events with different amplitudes in many recording channels. Hence, ABVA used the data of each shank to perform the spike detection and discrimination steps.

In the first stage of the algorithm, spikes are detected using a double threshold methodology, with a maximum time window *t*_*w*_ and a minimum threshold voltage *V*_*t*_*h*. Specifically, spikes are detected as spatiotemporally connected events coming from one cell when the snippets for different recording sites exceed the *V*_*t*_*h* and showed peak times below *t*_*w*_. The next step is feature extraction which obtains the d spike properties that allows larger spike discrimination. We used PCA to extract the most representative waveforms within the eight recording sites of a shank. So, for a spike consisting of n sample points (62 samples, corresponding to 2.5 ms) for each recording site, the feature extraction method produces m variables (m < n), where m is the number of features in PCA space (the number of features is a user-settable parameter, with default value 3).

The next step is spike clustering. In this stage spikes with similar feature vectors across the eight recording sites are grouped into clusters in the low-dimensional space, assuming that each cluster represents the spikes of a single cell. We use K-means clustering to classify the spatiotemporally connected events. The optimal number of clusters was evaluated by the MATLAB Statistics Toolbox function “evalclusters” based on the Calinski-Harabasz clustering criterion. Crucially, we built a mask that corresponds to the voltage shape templates of temporally overlapping events in the group of channels that were clustered as an individual cell (Figure S9A). This spike-shape template mask was used to discriminate during task performance the response events of a cell across the eight recording sites of a shank.

Finally, the results are curated to adjust the potential errors made by the clustering algorithm. In our case, the curation is made by the algorithm automatically, polishing the clustering results using a dual-threshold on the spike mask. This threshold is applied to each recording site template of the mask and includes a minimum number of occurrences as well as a minimum amplitude of the peak-to-valley waveform. This strategy strongly avoids spurious detection of small noise events on the masks. As a result, in one session, up to 142 isolated single neurons were detected (average of 102, cells per recording, range: 42-142), for 8 analyzed recording sessions in monkey 1 and 14 in monkey 2. The total number of recorded neurons was 1189, 225 for monkey 1 and 964 for monkey 2.

Our goal was to produce a practical system than can be used in our laboratory for processing information using high-count silicon probes offline and in a real-time way. We validated our algorithm by comparing ABVA with the commonly used KiloSort (Pachitariu et al., 2016). KiloSort (KS) 2.5 was used for spike discrimination, using all default parameters with the exception of ops.Th = [10,4]. We compared the results of both algorithms for different recording sessions using the Pearson correlation over the activation profiles of the resulting cell responses during the task performance of our four conditions. We identified the cells with high correlations (r2 > 0.3, P < 0.001) in their activity pattern between the two algorithms. Supplementary Figure S9C shows activation profiles of the 72 cells of session 1 of Monkey 2 with high correlations between algorithms, where KiloSort identified 138 and ABVA 99 total cells. Similar robust correlations in activity patterns were obtained between the two methods across all sessions.

### Neural activation periods

We used the Poisson-train analysis to identify the cell activation periods within each interval defined by two subsequent taps. This analysis determines how improbable it is that the number of action potentials within a specific condition (i.e. target interval and ordinal sequence) was a chance occurrence. For this purpose, the actual number of spikes within a time window was compared with the number of spikes predicted by the Poisson distribution derived from the mean discharge rate during the entire recording of the cell. The measure of improbability was the surprise index (SI) defined as:

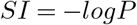

where P was defined by the Poisson equation:

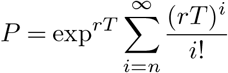

where P is the probability that, given the average discharge rate r, the spike train for a produced interval T contains n or more spikes in a trial. Thus, a large SI indicates a low probability that a specific elevation in activity was a chance occurrence. This analysis assumes that an activation period is statistically different from the average discharge rate r, considering that the firing of the cell is following a non-homogenous Poisson process (Perez et al., 2013). The detection of activation periods above randomness has been described previously (Merchant et al., 2001, 2015a) (Merchant et al., 2001; Merchant, Pérez, et al., 2015). Importantly, the Poisson-train analysis provided the response-onset latency and the activation period for each cell and for each combination of target interval/serial order.

### Neural trajectories

For each recording session and for each neuron, we calculated the produced interval (time between two taps) across repetitions, and it was divided off into a variable number of bins *Bin*_*size*_, this number depended on the target interval of the trial, it is called *Bin*_*time*_. For example, the total number of bins was 22 and 42 for the target intervals trials of 450 and 850 ms, respectively. This is called the target interval of normalized data (TIND):

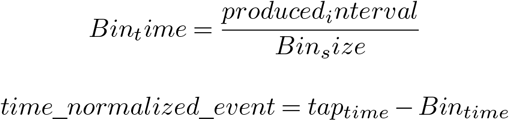

We binarized the neural data by calculating the discharge rate on each bin. After that, we computed the average firing rate in non-overlapping time bins smoothed with a Gaussian kernel (sigma = 20 ms). Finally, the binarized data of each neuron was normalized.

### Principal component coefficients matrix

Given a linear transformation of a matrix *X* into a matrix *Y*, such that each dimension of *Y* explains variance of the original data *X* in descending order, PCA can be described as the search for matrix *P* that transforms *X* into *Y* as follows:

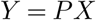

Hence, we first calculated the matrix *P* using a matrix *X* that includes all trials and target interval combinations for the visual and auditory ST of our TIND cell population. Using this P on other data guarantees that the same transformation is applied to different neural activity sets. Therefore, using the TIND framework we avoided over-or under-representation of the information for different target intervals, due to the constant total number of bins across conditions.

### Gaussian Process Factor Analysis

Gaussian Process Factor Analysis (GPFA) extract low-dimension latent trajectories from noisy, high-dimension time series data (Yu et al., 2009). It combines linear dimensionality reduction (factor analysis) with Gaussian-process temporal smoothing in a unified probabilistic framework.

The input consists of a set of trials (*Y*), that includes all trials and target interval combinations for the visual and auditory ST of our TIND cell population, each containing a list of spike trains (Ν neurons). The output is the projection (*X*) of the data in a space of pre-chosen dimensionality *x*_*dim*_ < *N*.

Under the assumption of a linear relation (transform matrix *C*) between the latent variable *X* following a Gaussian process and the spike train data Y with a bias d and a noise term of zero mean and (co)variance *R* (i.e., *Y* = *CX* +*d*+ *N* (0, *R*)), the projection corresponds to the conditional probability *E*[*X* | *Y*]. The parameters (*C, d, R*) as well as the time scales and variances of the Gaussian process are estimated from the data using an expectation-maximization(EM) algorithm.

### Generating neural trajectories

The TIND information for every trial of all neurons constituted the columns of the matrix. The principal component coefficients matrix P were multiplied by the *X*^*′*^ matrix to transform the neural data into the space of the original *Y*. Using the same transformation matrix for each trial allowed the comparison of trajectories for different trials and tasks. A locally weighted scatterplot smoothing function was applied to the columns of the *Y* matrix. The first three dimensions of Y were used to generate graphical three-dimensional trajectories.

### Coding subspaces and mixed variance

To identify coding subspaces, we decomposed neural population activity based on the covariance of the neural activity for each of the following parameters: Duration, Modality, Elapsed Time, and Tapping Times. To define the 3D coordinate system of these subspaces, we calculated the covariance matrices onto the projected neural population activity (first three PCs) and compute the eigen values as follows:

We used the projected data (*Y*) onto the first three PCs, which includes all trials and target interval combinations for the visual and auditory ST. Duration parameter was arranged in a matrix *R*^(^[8400*x*3]), with each column associated to each PCs and rows corresponding to the averaged neural state for all trials and modalities along two durations. In the case of the modality parameter, the matrix includes the averaging along the two modalities instead of durations. Finally, the coding subspace of Elapsed time parameter was arranged in a matrix *R*^(^[4200*x*3]), with each column associated to each PCs and rows corresponding to the averaged neural state along two durations and two modalities of all trials and target interval. To calculate this subspace, we used the sum of the diagonal elements of the covariance matrix.

### Trajectory amplitude, angle, and position

The first three PCs explained the 7.1, 4.1 and 4 percent of the total variance. The PC1 showed a steep change at the beginning and end of the trial, suggesting a chunking mechanism of the tap sequence in the overall of the neural population-state. In contrast, the PC2 and PC3 showed a strong oscillatory structure with a phase difference of 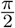 radians during SC. For these three PCs, we calculated the amplitude of the trajectory as the mean of the Euclidean distances between the anchor point (Figure 2C) and each point in the trajectory segment across the four serial order elements for each target interval.

The angle of the trajectory was calculated as the mean of the angle from dot product between the anchor point (Figure 2C) and each point in the trajectory segment across the four serial order elements for each target interval. The variability of position was calculated as the standard deviation of the distances between the anchor point (Figure 2D) and each point in the trajectory segment across the four serial order elements for each target interval.

### Movement kinematics

We applied the Lucas-Kanade optic flow method to measure the monkey’s arm speed during the ST. This method calculates a flow field from the intensity changes between two consecutive video frames. The analyzed video was recorder with a high-speed camera (Basler acA750 AG) positioned orthogonally to the hand’s plane of motion with a 640×480 resolution at 250 frames per second. The optic flow method was applied to a smaller area of 140×140 pixels from the original video that contained the monkey’s arm during the whole trial and no other moving objects. The arm’s movement velocity vector was calculated across all frames as the magnitude of the sum of all the individual flow fields vectors whose magnitude was larger than a predefined threshold. The velocity vector was calculated from the first to the last tap on each correct trial. We reported the speed as the magnitude of the velocity vector (Barron et al., 1994). Posteriorly, the kinematic state of the arm was tagged as movement when the velocity vector was larger than a threshold or dwell otherwise. The tagging algorithm considered a change on the kinematic state when the new state lasted longer than 3 consecutive frames.

### Temporal scaling in the synchronization task

To quantify temporal scaling, we defined the scaling index (SI) of a subspace S as the portion of variance of the projections of trajectories into S that can be explained by temporal scaling (Wang et al., 2018). We computed the kth scaling component *u*_*SC,k*_ as:

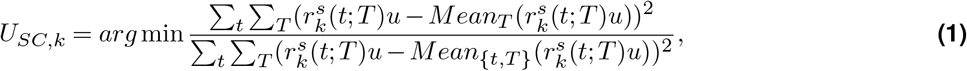

where 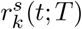 is population activity at the scaled time when the duration of the production epoch is T, the denominator is the total variance of the trajectories, and the numerator is the variance that cannot be explained by temporal scaling. As in (Bi & Zhou, 2020) and (Zhou et al., 2022), for computing the first scaling component *U*(*SC*, 1) we set 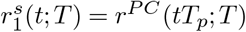 with 0 ≤ *t* ≤ 1, where *r*^*P C*^ is the projection of the population activity in the subspace spanned by the first 3 PCs, and *T*_*p*_ is the produced interval by the monkey in the production epoch; then we minimized*−*u in eq. 1. To calculate the second scaling component *U*(*SC*, 2) we set 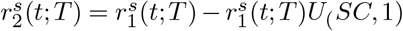, and then minimized u in eq. 1 in the subspace orthogonal to *U*(*SC*, 1). In this way, we computed all the 3 scaling components one at the time. Finally, the scaling index (SI) of a subspace was defined as:

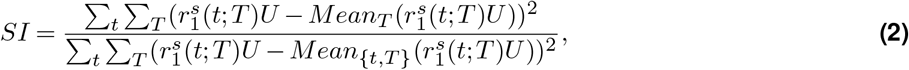

where 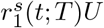 is the projection of the scaled trajectory to the subspace *U*. For implementation we used https://github.com/zedongbi/IntervalTiming (Bi & Zhou, 2020).

### Moving bumps simulations

In order to investigate how the properties of neural sequences were associated with the geometry and kinematics of population neuronal trajectories we performed different simulations. Each simulation comprises a set of *N*_*d*_ neurons with a firing rate:

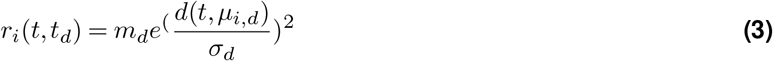

where t is the elapsed time, *m*_*d*_ and *σ*_*d*_ are maximum discharge rate and duration of the activation period, respectively, which depend on the target duration *t*_*d*_.*µ*_*i,d*_ corresponds to the time of peak activation of neuron *i*. The peaks of activity at *µ*_*i,d*_ cover the entire interval *t*_*d*_. In order to create neural sequences that reset at the next produced interval in the sequence used the distance function *d*(*t, µ*_*i,d*_) that is congruent with the difference *t µ*_*i,d*_modulo *t*_*d*_, allowing for the generation of cyclic kinematics.

Regarding the duration of cell activity, for figures 7C,E and F we used *σ*_8_50 *> σ*_4_50 or *σ*_8_50 = *σ*_4_50 for the remaining simulations. For the response magnitude, in Figure 7D we used *m*_8_50 *> m*_4_50 and *m*_8_50 = *m*_4_50 for all the other cases. The dynamics of the activation peak of Figures 7A, 7B, and the shared cells at the beginning and end on the interval of Figures 7E-G, followed *µ*_*i*,450_ = *µ*_*i*,850_, for the remaining simulations 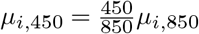. Finally, we used *N*_4_50 *< N*_8_50 for figures 7B and F. For Figures 7F and G we inserted randomly neurons within the intermediate portion of the produced interval.

## Acknowledgments

We thank Ranulfo Romo, Pavel Rueda, Jonathan Cannon, and Roman Rossi for their fruitful comments on the manuscript. We also thank Raul Paulín and Luis Prado for their technical assistance.

Abraham Azahel Betancourt Vera is a doctoral student from the Programa de Doctorado en Ciencias Biomédicas, Universidad Nacional Autónoma de México (UNAM) and has received CONACYT fellowship 403878.

Hugo Merchant is supported by Consejo Nacional de Ciencia y Tecnología (CONACYT) Grant CONACYT: A1-S-8430, UNAM-DGAPA-PAPIIT IN201721.

