## Supplemental figures and tables for "Amodal population clock in the primate medial premotor system for rhythmic tapping"

### Supplementary Information

| | Movement time | | | | Autocorrelation Lag-1 PI | | | | |
|---|---|---|---|---|---|---|---|---|---|
| #Session | Movement amplitude | | Movement speed | | Autocorrelation Lag-1 Mov. amplitude | | Autocorrelation Lag-1 Mov. speed | | #Cells |
| | r | p | r | p | r | p | r | p | |
| 2 | 0.17 | 0.21 | −0.02 | 0.86 | 0.26 | 0.01 | 0.26 | 0.01 | 42 |
| 3 | −0.03 | 0.81 | −0.27 | 0.05 | 0.03 | 0.75 | 0.03 | 0.75 | 28 |
| 6 | −0.45 | 0.0001 | −0.51 | 0.0001 | 0.1 | 0.3 | 0.1 | 0.3 | 97 |
| 8 | −0.04 | 0.79 | 0.33 | 0.01 | −0.02 | 0.87 | −0.02 | 0.87 | 76 |
| 9 | −0.61 | 0.0001 | −0.17 | 0.12 | 0.18 | 0.08 | 0.18 | 0.08 | 84 |
| 11 | −0.4 | 0.0001 | 0.14 | 0.28 | −0.09 | 0.36 | −0.09 | 0.36 | 75 |
| 14 | −0.16 | 0.28 | −0.16 | 0.26 | 0.08 | 0.41 | 0.08 | 0.41 | 21 |
| 1 | 0.23 | 0.05 | −0.24 | 0.04 | 0.12 | 0.23 | 0.12 | 0.23 | 92 |
| 4 | −0.27 | 0.04 | −0.03 | 0.84 | 0 | 1 | 0 | 1 | 97 |
| 5 | −0.4 | 0.0001 | −0.11 | 0.4 | 0.19 | 0.06 | 0.19 | 0.06 | 91 |
| 7 | −0.17 | 0.22 | 0.39 | 0.0001 | −0.16 | 0.11 | −0.16 | 0.11 | 94 |
| 10 | 0.06 | 0.67 | −0.62 | 0.0001 | 0.21 | 0.03 | 0.21 | 0.03 | 81 |
| 12 | −0.06 | 0.0001 | −0.27 | 0.07 | 0 | 0.96 | 0 | 0.96 | 64 |
| 13 | 0.2 | 0.15 | 0.02 | 0.9 | 0.02 | 0.81 | −0.02 | 0.81 | 22 |
| Significant sessions | 1 | | 2 | | 2 | | 2 | | |

**Table S1.** Individual session analysis where we computed the Pearson correlation coefficient between the behavioral (top labels) and kinematic parameters of the neural trajectories (bottom labels) in the fourteen sessions of Monkey 2 with collected videos on behavior. Orange and blue shading follows the significant effects of Table 1.

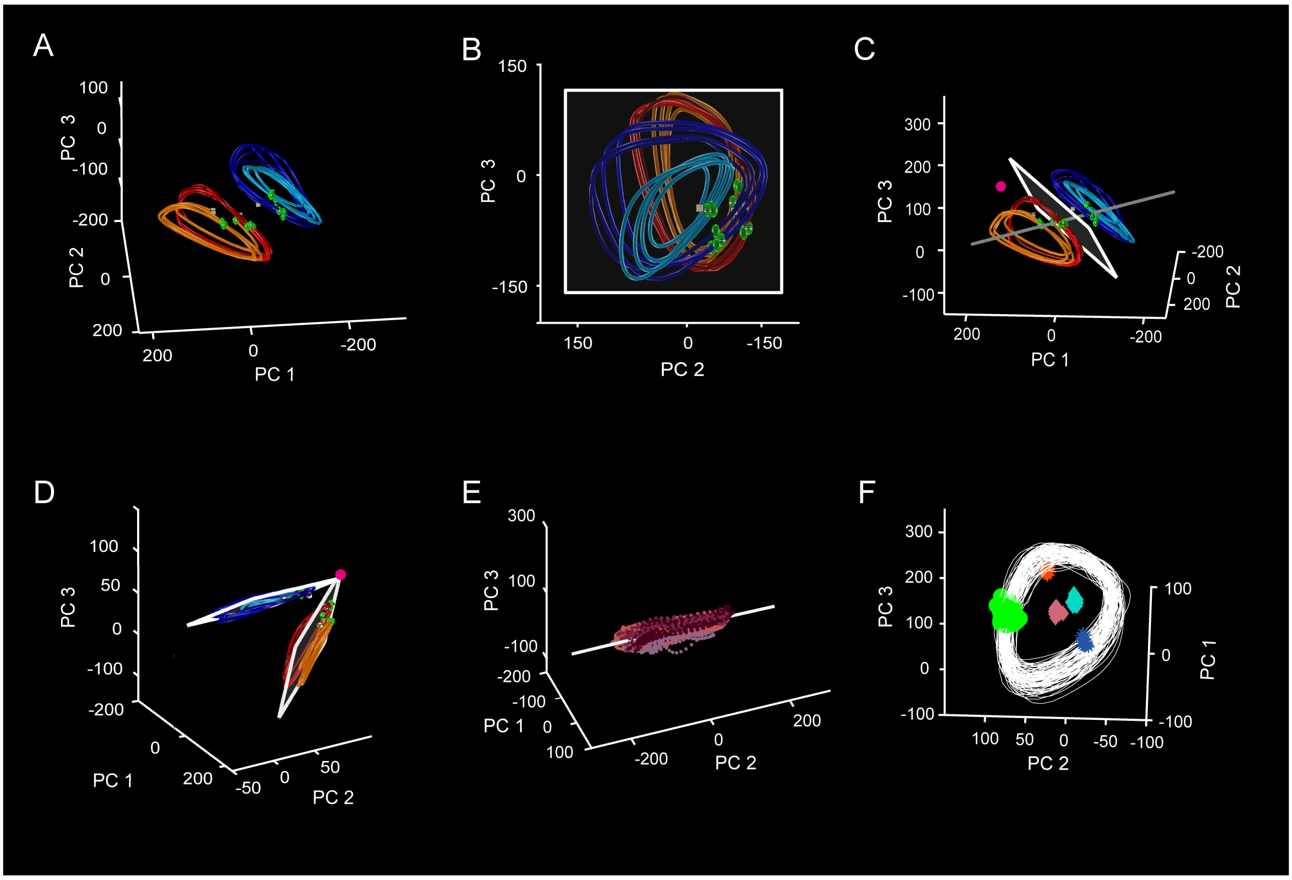


**Figure S1. Neural population trajectories during ST and their oscillatory dynamic properties using alternative reduction method (GPFA). Format as in Figure 2.**

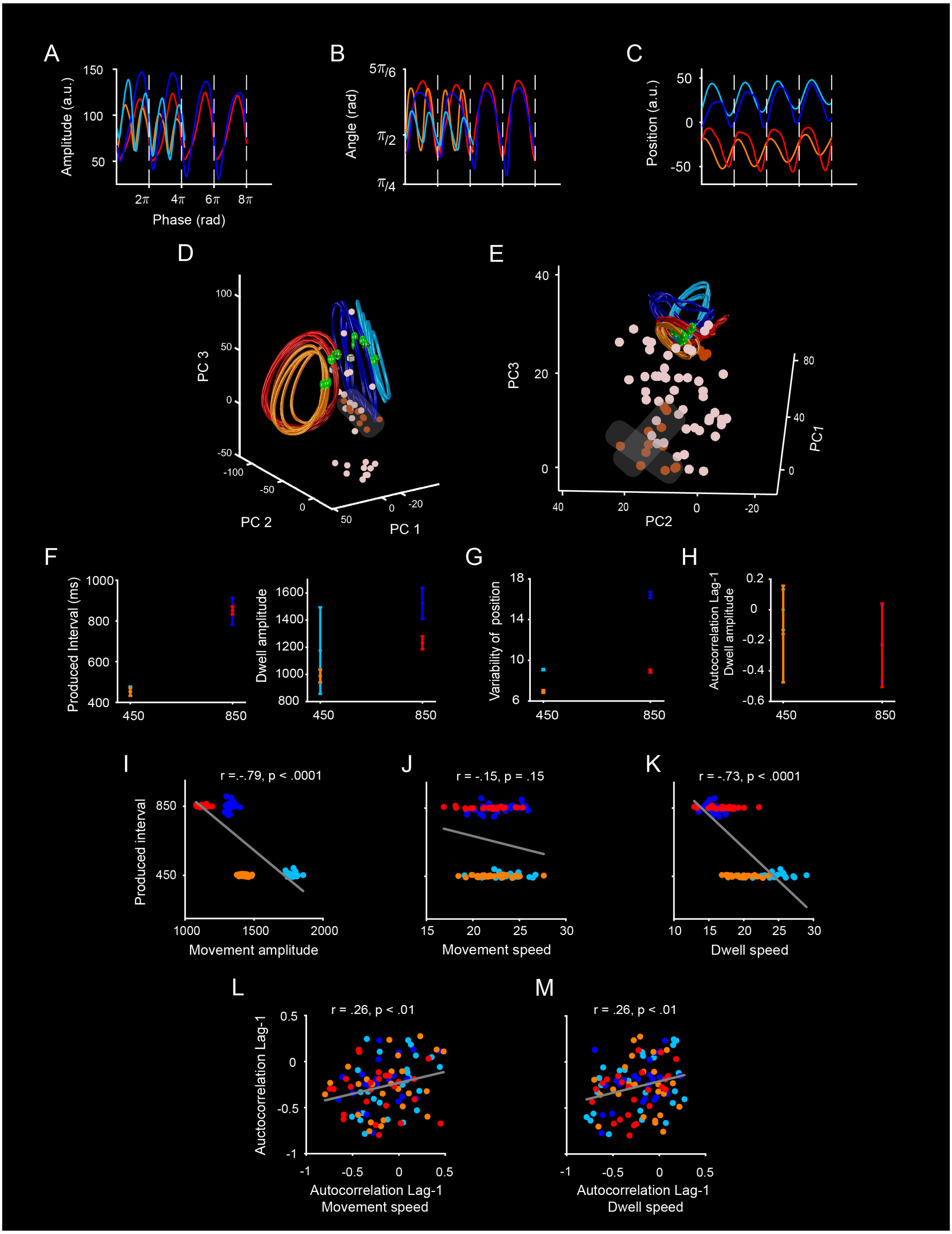


**Figure S2. Kinematic of neural trajectories in stable points from a distribution of arbitrary points onto the state space.**

**Figure S2 *(previous page)*.** (A) Neural trajectories in the first three PCs as in Figure 2A. The tested arbitrary points for computing the amplitude and angle are depicted in state space with two colors. Dark orange for the positions where the properties of behavior were replicated in the neural population kinematics, as in Figure 3A-C and pink for the positions where the kinematics did not follow these properties. Note that similar kinematic properties were obtained when the arbitrary point was located within a contiguous region (shaded) in state space.

(B) The same as in A but only for the projected neural activity of Session 2 of Monkey 2.

(C) Dwell and Movement time amplitude (± 2xSEM), computed as area under curve from neural trajectories of Session 2 of Monkey 2, as a function of target interval. The ANOVA showed significant main effects of modality, $F(1,196) = 1580.9$, $p < .0001$; epoch, $F(1,196) = 3716.2$ , $p < .0001$; but not statistical significance on duration, $F(1,196) = .45$, $p = .5$; as well as significant effects on duration x epoch interaction, $F(1,192) = 4194.82$, $p < .0001$ and modality x epoch interaction, $F(1,196) = 38.7$, $p < .0001$.

(D) Lag 1 autocorrelation of the amplitude of the neural trajectories during the Dwell time as a function of target duration for Session 2 of Monkey 2. The ANOVA showed significant main effects of duration, $F(1,196) = 4.17$, $p < .04$, but no statistical significant effect on modality, $F(1,196) = 3.21$, $p = .07$ and on duration x modality interaction, $F(1,196) = .14$, $p = .70$.

(E) Variability of the position (SD within and across trials) as a function of target interval (± 5xSEM) for the recording session in D. The ANOVA showed significant main effects of duration, $F(1,196) = 26.1$, $p < .0001$, modality, $F(1,196) = 31.63$, $p < .0001$; as well as significant effects on duration x modality interaction, $F(1,196) = 20.3$, $p < .0001$.

(F) Negative significant correlation between the produced interval and movement amplitude ($r = -0.79$, $p < 0.0001$) for the recording session in D.

(G) Not significant correlation between the produced interval and movement time speed of the neural trajectories ($r = -0.15$, $p = 0.14$) for the recording session in D.

(H) Negative significant correlation between the produced interval and dwell time speed of the neural trajectories ($r = -0.73$, $p < 0.0001$) for the recording session in D.

(I) Significant correlation between the autocorrelation Lag-1 of the produced interval vs autocorrelation Lag-1 of the movement amplitude ($r = 0.26$, $p < .007$) for the recording session in D.

(J) Significant correlation between the autocorrelation Lag-1 of the produced interval vs autocorrelation Lag-1 of the movement speed ($r = 0.26$, $p < .007$) for the recording session in D.

(K) Significant correlation between the autocorrelation Lag-1 of the produced interval vs autocorrelation Lag-1 of the movement speed ($r = 0.25$, $p < .009$) for the recording session in D.

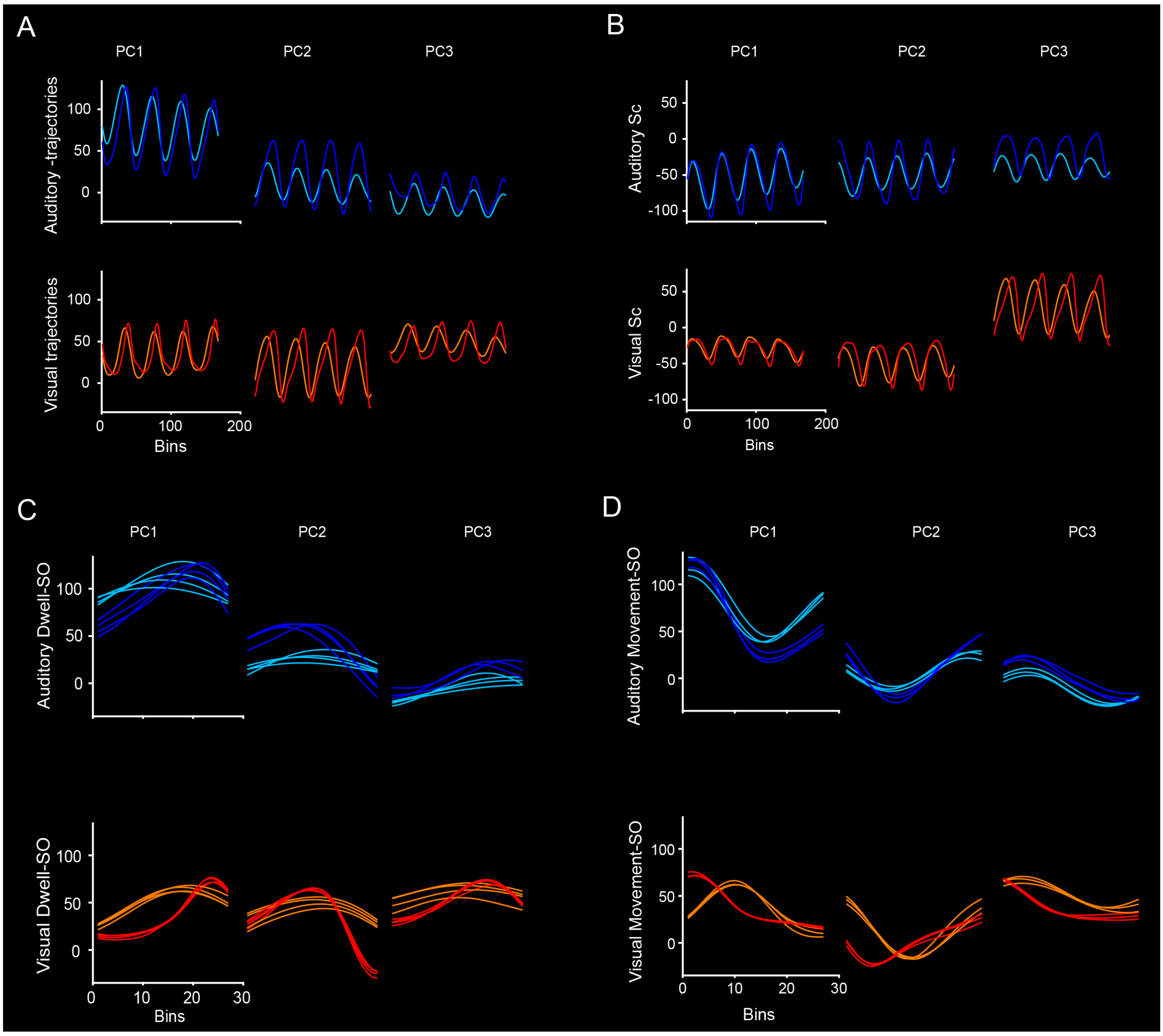


**Figure S3. Neural population traces.**

(A) Neural population geometry of the first three principal components of the auditory and visual modality used to calculate the scaling index and scaling components. Color code format as in Figure 2B. (B). Scaling components for the auditory (Top panels) and visual (Bottom panels) modality calculated in the subspace spanned by the first three principal components.

(C) Portion of the 3 PCs during dwell time for the auditory (top) and visual (bottom) modalities.

(D) Same es in C but for the movement time epoch of the ST.

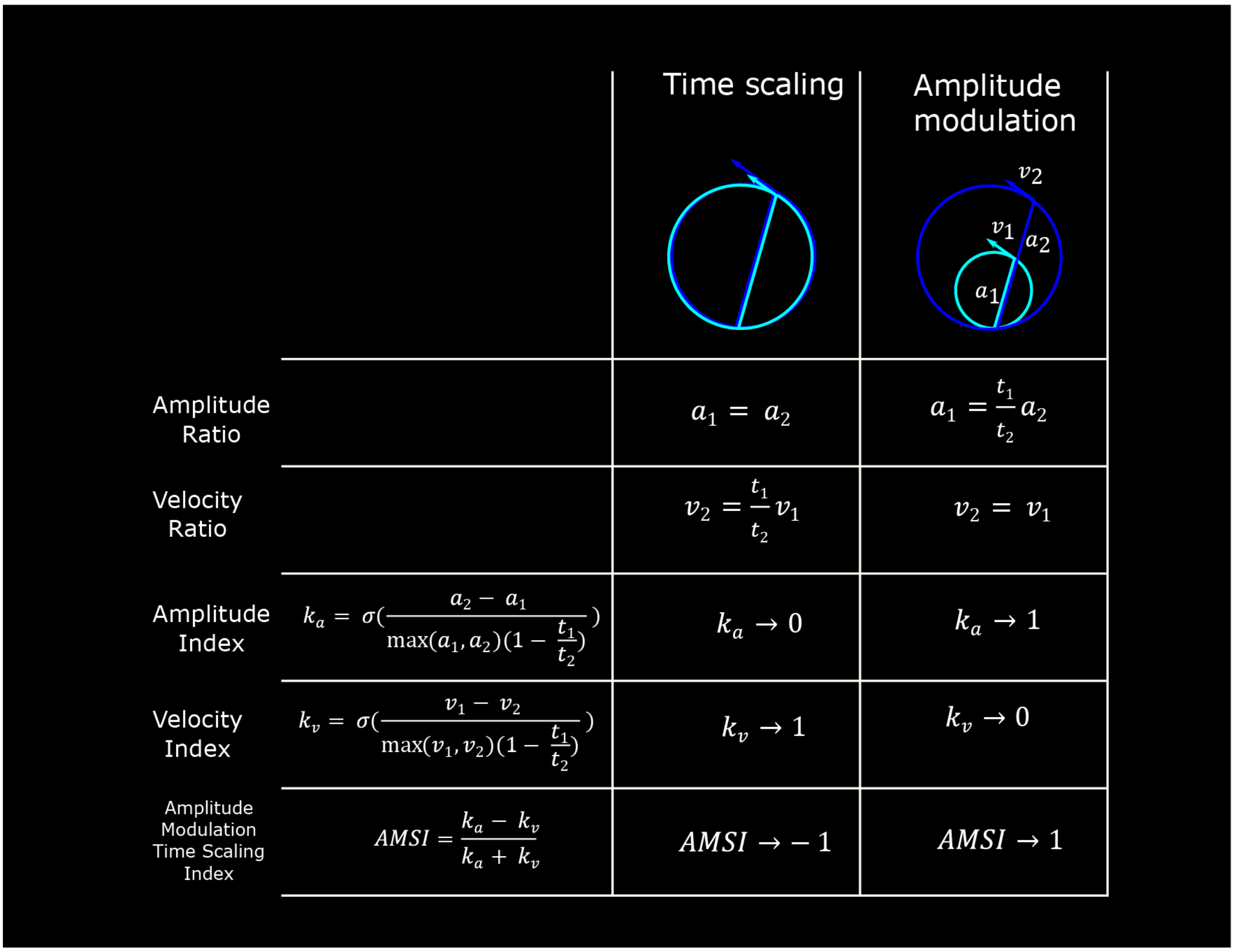


**Figure S4. Amplitude-modulation time-scaling index (AMSI).**
Top. Geometric description of the changes in amplitude (a) and velocity (v) for trajectories that show a full temporal scaling (Velocity Index = 1; AMSI = -1) or a full amplitude modulation (Amplitude index = 1; AMSI = 1). t1 = 450, t2 = 850 ms.
Bottom. Equations for all velocity and amplitude ratios and indexes.

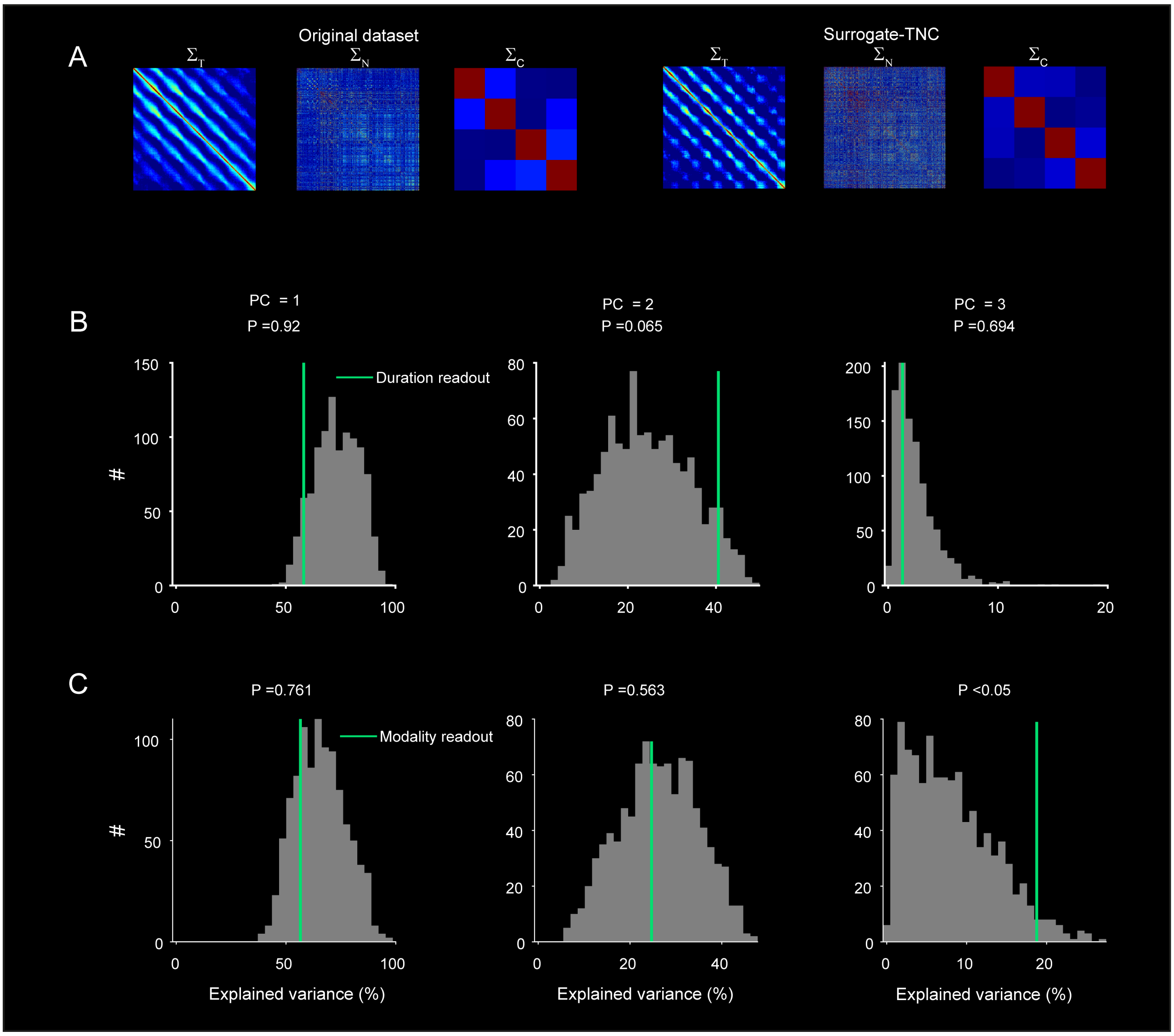


**Figure S5. Tensor Maximum Entropy (TME) surrogate preserve the specified primary features (TNC).**
(A) Heatmaps show the three covariances matrices of the original population neural responses across times ($\Sigma_T$), neurons ($\Sigma_N$), and conditions ($\Sigma_C$) using TME.
(B) Distribution of variance-explained on the top 3 principal components (PCs) from 1000 surrogate datasets from duration neural population. Malachite vertical lines mark value for the real explained variance.
(C) Distribution of variance-explained on the top 3 principal components (PCs) from 1000 surrogate dataset from modality neural population. Malachite vertical lines mark value for the real explained variance.

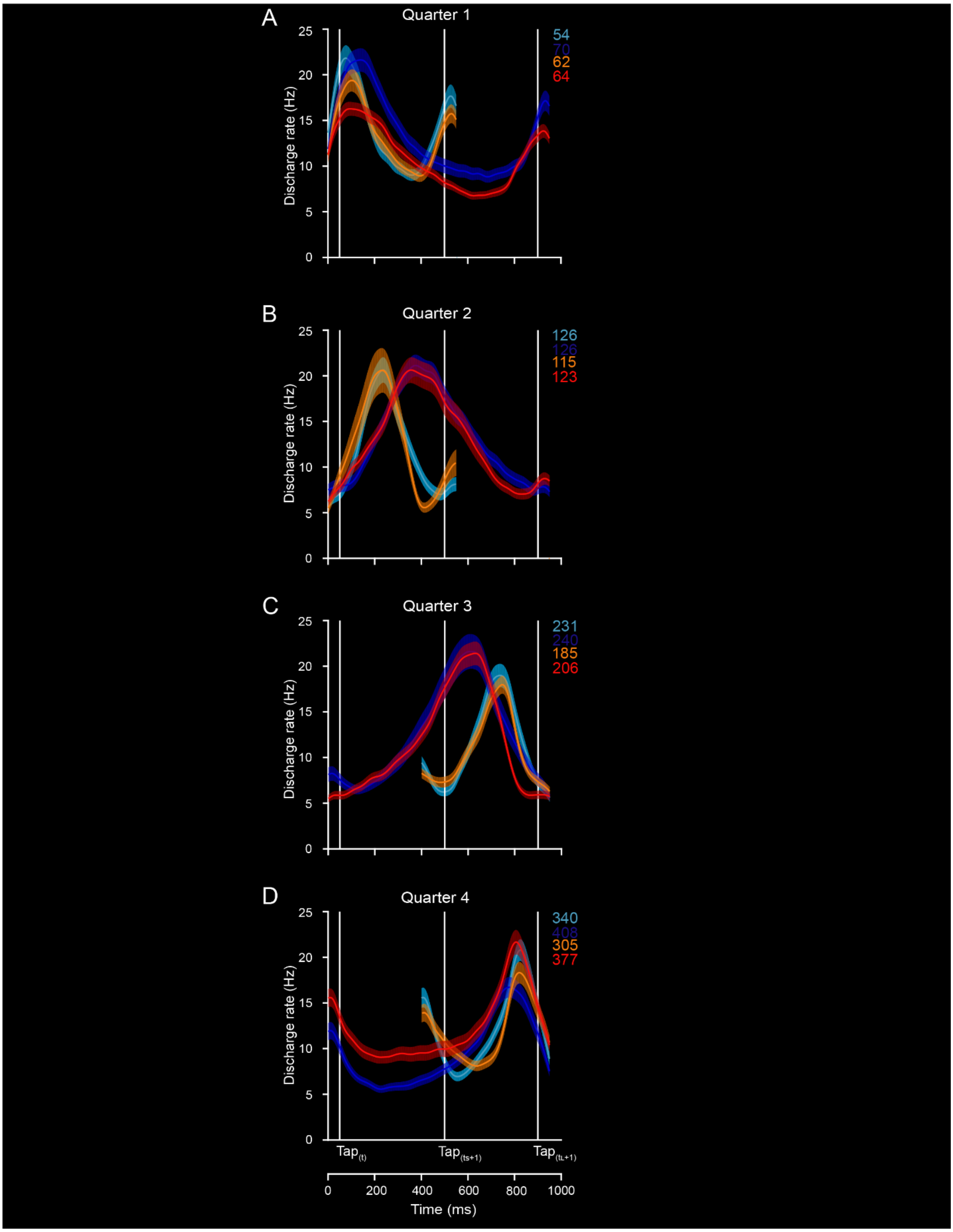


**Figure S6. Population SDF (mean ± SEM) of the neurons with activation periods during each of the four quarters of a produced interval. The neurons of moving bumps showed instantaneous activity changes that correspond to different types of ramping patterns reported previously.**

(A) Neurons of the first quarter showed ramping profile of the swinging ramps.

(B-C) The neurons of the second and third quarters showed an up-and-down profile of activation that correspond to ramps that encode elapsed time in the width or height of the ramp.(D) In the last quarter, the neurons encode the time-remaining for an action, reaching a peak at a particular time before the next tap.

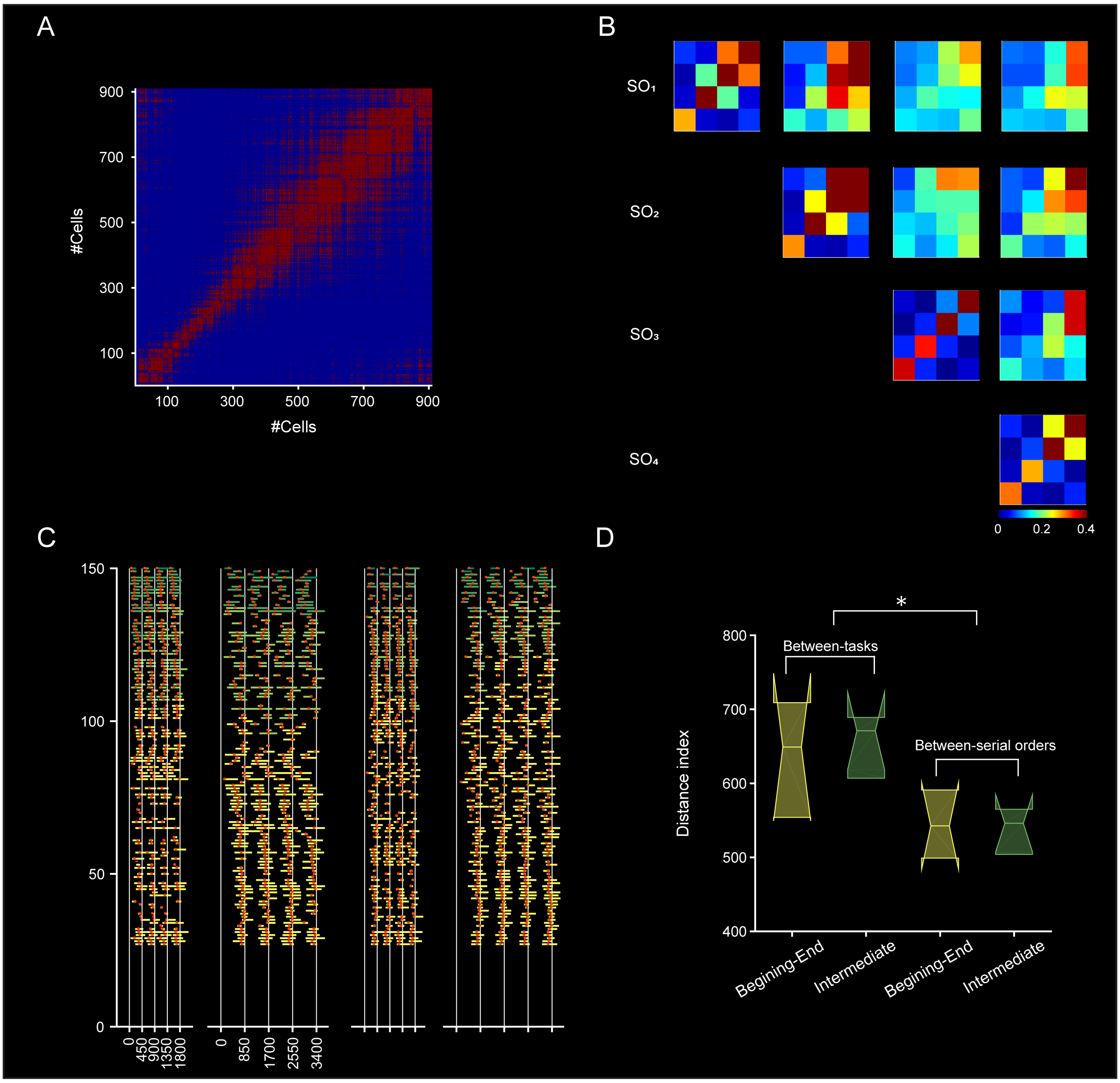


**Figure S7. Activation profiles across the four conditions and the four serial order elements of the ST.**

(A) Correlation matrix of self-sorted cells for auditory short condition (significant r-squared correlations between pairs of cells are illustrated as red points). The significant correlations were distributed not only around the diagonal but also for pairs of cells that were active at the beginning and end of the neural sequence.

(B) Probability that pairs of cells showed large correlation values ($r > 0.5$) in their response profile dividing the complete neural sequence into quarters. The color code goes from zero for blue quarter to one for red quarters. Large probability values are manly observed for the last two quarters of the neural sequences across all pairwise task comparisons. (C) Activation periods for the 124 neurons that showed similar response profiles across the four conditions and the four serial order elements of the ST. Figure conventions as in Figure 4A. (D) Distance index for initial intermediate and final bins using the Euclidean distance across task conditions (left, from Figure 6B) and serial order elements of the auditory large condition (right, from Figure 6C).

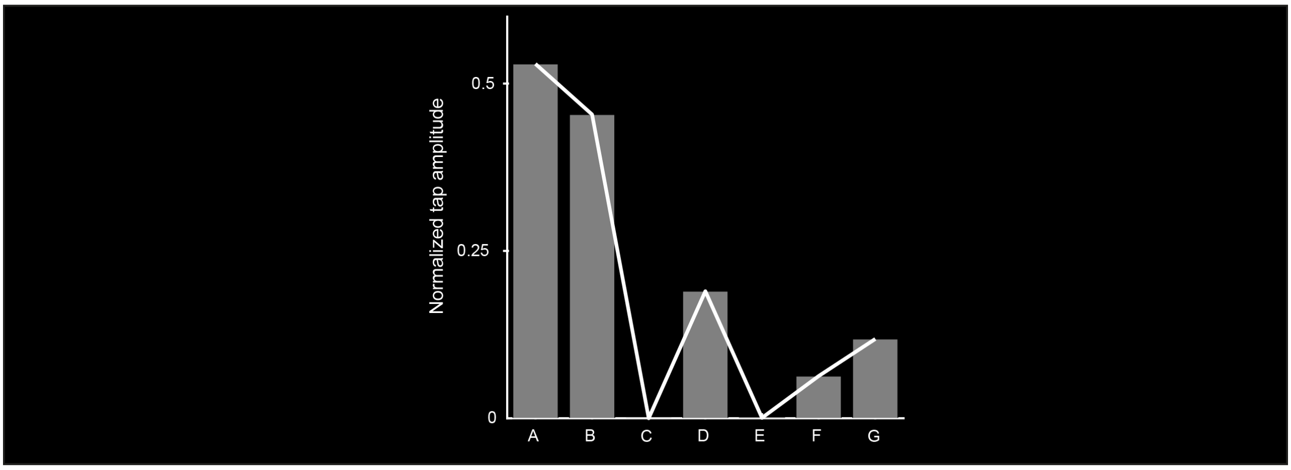


**Figure S8. Normalized distance between the short and long interval trajectories at the tapping times for the simulations in Figure 7 panels A-G**

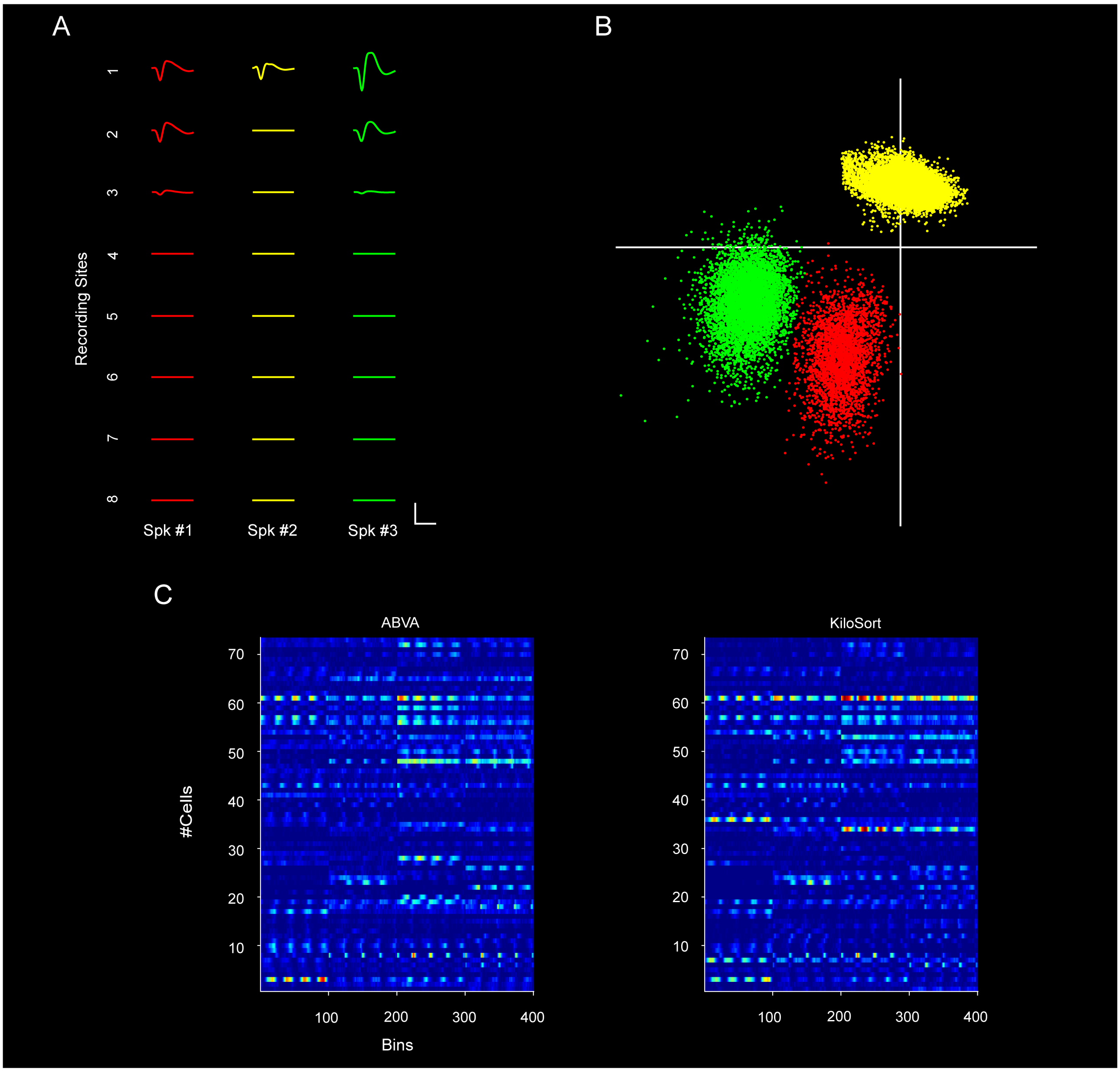


**Figure S9. Semiautomatic spike discrimination of the cells with ABVA.**

(A) Waveforms (mean ± SD) of the 4-clustered cells (Spk1-Spk3) across the 8 neighboring recording sites of 1 silicon probe with eight recording sites.

(B) Spike clusters of the three cells displayed along the first principal components of spike waveforms extracted from the 1st, 3rd, 4th and 6th channel. Same color code as in A. C: calibration: vertical 150 µV; horizontal 2 ms.

(C). Cells showed large numbers of neurons with significant correlations in their response profiles between spike sorting frameworks for one recorded session. We show the discharge rate of neural activity sorted with ABVA (left panel) and KiloSort (right panel). We found 72 cells with a high correlation ($r > .3$) in their response between sorting methods ($p < .0001$).
